# Old Yellow Enzyme from *Brevibacillus nitrificans* functions as 12-*oxo*-phytodienoic acid reductase *in planta*

**DOI:** 10.64898/2026.05.27.728186

**Authors:** Moritz Klein, Ellen Hornung, Liv Perle, Kirstin Feussner, Cornelia Herrfurth, Alisa Keyl, Lars Bröker, Lasse Stöhr, Stefan A. Rensing, Mats Hamberg, Jan de Vries, Ivo Feussner

**Author notes:** **Corresponding author** I. Feussner; Dept. for Plant Biochemistry, Albrecht-von-Haller-Institute for Plant Sciences, University of Goettingen, Justus-von-Liebig-Weg 11, 37077 Goettingen, Germany. **Email addresses:** Moritz Klein,; Ellen Hornung; Liv Perle;, Kirstin Feussner; Cornelia Herrfurth; Alisa Keyl; Lars Bröker; Lasse Stöhr;, Stefan A. Rensing; Mats Hamberg; Jan de Vries.

## Abstract

Old Yellow Enzymes (OYEs) are a widely distributed family of *ene*-reductases that were first described in a *Saccharomyces cerevisiae* ferment. In plants, *cis*-12-*oxo*-phytodienoic acid (*cis*-OPDA) reductase (OPR) is the best studied OYE. In *Arabidopsis thaliana*, the peroxisomal AtOPR3 was characterized as the major OPDA reductase, which generates 3-*oxo*-2-(2-pentenyl)-cyclopentane-1-octanoic acid in the jasmonic acid (JA) biosynthesis. In *Atopr3* lines, only small amounts of JA are detectable after wounding. Here, we describe an OPR-like enzyme (named BnOPR) from the gram-positive *Brevibacillus nitrificans*. The sequence was identified in an early version of the *Physcomitrium patens* genome and is assumed to be a contamination by a bacterium growing in association with *P. patens*. In complementation experiments with an *Atopr3* line, we demonstrate that expression of BnOPR, fused with a peroxisomal targeting signal, rescues the male infertile phenotype and increases JA and JA-Ile levels. The catalytic parameters of BnOPR were determined for a set of substrates, including *cis*-OPDA and prednisone. Interestingly, *B. nitrificans, B. brevis*, and *Paenibacillus physcomitrellae* were shown to have a positive effect on *P. patens* growth.

**Highlight:** The bacterial enzyme BnOPR rescues the male infertile phenotype of *Atopr3* plants.

## Introduction

Old Yellow Enzymes (OYEs; EC 1.6.99.1) are a family of reductases acting on α,β-unsaturated carbonyl compounds by the addition of a proton and a hydride ion. The class was first described in *Saccharomyces carlsbergensis*, which is today referred to as *S. pastorianus*, as OYE and later OYE2 (Saito *et al*., 1991; Stott *et al*., 1993; Warburg and Christian, 1932). OYEs share a common TIM-barrel fold with eight α-helices surrounding the eight-stranded β-barrel (Fox and Karplus, 1994). Chromatographic analysis revealed the interaction of two or more OYE to form homo- or heterodimers. Further extensive research led to an expanding enzymatic knowledge of OYE in bacteria and plants (Li *et al*., 2009; Toogood *et al*., 2010). While numerous candidates were investigated towards their application in chemical synthesis and bioconversion of reactive substances, the *in vivo* activity and function of most OYEs is still unknown. The most studied OYE with a confirmed specific *in vivo* reaction is the plant *cis*-12-*oxo*-phytodienoic acid (*cis*-OPDA) reductase (OPR) involved in jasmonic acid (JA) biosynthesis (Schaller *et al*., 2000; Stintzi and Browse, 2000). Lack of OPR activity in *Arabidopsis thaliana* leads to an impairment inanther development and pollen germination, and mutant plants are considered to be male sterile (Stintzi and Browse, 2000). This phenotype could be rescued by applying methyl jasmonate (MeJA). As expected, OPR3 deficient plants produced no JA upon wounding (Stintzi *et al*., 2001). However, the defense response was not completely defective as *Atopr3-1* plants could resist insect feeding and fungal pathogen attacks with a wild-type-like survival rate. Even though, reassessment of the *Atopr3-1* mutant plants reported a conditional production of JA, the speculations about another isoform that could reduce cyclopentenones *in planta* and produce the bioactive JA stereoisomer were confirmed when AtOPR2 was identified as a reductase for the cyclopentenone 4,5-ddh-JA (Chehab *et al*., 2011; Chini *et al*., 2018). *Atopr3-3 Atopr2-1* plants were nearly completely deficient in JA and showed a decreased transcriptional response to *Botrytis cinerea* infections compared to the single mutant, *Atopr3-3*. Similar observations were made in other plants such as tomato and corn (Strassner *et al*., 2002; Yan *et al*., 2012).

Besides these plant OPRs, several bacterial and fungal OYEs were characterized towards potential *in vivo* functions and *in vitro* substrate specificity. For an expanding number of OYEs, a classification system based on phylogenetic relations was proposed with class I to class V (Peters *et al*., 2019). Class I is divided into the subclasses Ia of pentaerythritol tetranitrate reductase (PETNR)-like OYEs, Ib of OPR-like OYEs, and Ic with OYE1-like OYEs (Shi *et al*., 2020). Class II consists of the thermophilic-like OYE with yqjM from *Bacillus subtilis*, which shows an upregulation upon exposure to nitro compounds like 2,4,6-trinitrotoluene (TNT), nitroglycerin, and oxidative stress (Fitzpatrick *et al*., 2003). Furthermore, thermophilic “ene” reductases (TOYEs) show an oligomer formation by interaction of C-terminal residues and the active site (Adalbjörnsson *et al*., 2010). This differs from the reported self-inhibition mechanism by dimerization, which was described for tomato (*Lycopersicon esculentum*) LeOPR3 (Breithaupt *et al*., 2006). TOYEs show increased stability against higher temperatures and solvents, making them prime candidates as biocatalysts (Breithaupt *et al*., 2001). Class III and class IV consist of bacterial OYEs that show characteristic sequence similarities to the previous classes I and II but differ in substrate specificities and form clusters in phylogenetic analysis (Shi *et al*., 2020). Class V includes fungal OYEs. In a recent comprehensive study to expand the knowledge of OYEs, sequence similarity network analysis revealed further sub-clusters within these classes (White *et al*., 2025).

In the past, OYEs from the different classes were characterized in order to study their *in vitro* potential for chemical conversions with high stereoselectivity (Shi *et al*., 2020). Most studies focused on the classical OYEs from Class I, OYEs from *S. carlsbergensis*, and OPRs from corn, tomato, and *A. thaliana* (Breithaupt *et al*., 2001; Saito *et al*., 1991; Schaller *et al*., 2000; Schaller *et al*., 1998; Stott *et al*., 1993; Vick and Zimmerman, 1986). They bind phenolic compounds and reduce different quinones, like menadione and duroquinone, 2-cyclohexen-1-one, but also cyclopentenones like *cis*-OPDA (Fitzpatrick *et al*., 2003; Niino *et al*., 1995; Stott *et al*., 1993; Vaz *et al*., 1995). In a more recent approach, yqjM was used in combination with a squalene hopene cyclase from *Alicyclobacillus acidocaldarius* to convert citral to enantiopure (–)-*iso*-isopulegol (Peters and Buller, 2019). Another OYE, morphine reductase (MR), as a class I classical OYE from *Pseudomonas putida*, was found to reduce codeinone and morphinone *in vitro* (Barna *et al*., 2002; French *et al*., 1999). PETNR from *Enterobacter cloacae* PB2 and glycerol trinitrate reductase (NerA) from *Agrobacterium radiobacter* demonstrated the potential application in the reduction of nitro compounds, like the nitro-aromatic TNT (French *et al*., 1996; French *et al*., 1998; French *et al*., 1999; Oberdorfer *et al*., 2013). This finding expanded the circle of putative substrates and applications, as well as the potential of OYEs as detoxifying enzymes *in vivo*. *In vivo* studies with *A. thaliana* revealed the involvement of AtOPRs, predominantly AtOPR1 and AtOPR2, in the detoxification of TNT (Beynon *et al*., 2009). FOYE-1, from the acidophilic iron-oxidizing bacterium “Ferrovum” sp. JA12, showed high stability until temperatures up to 60 °C and showed the highest specific activity with (*R*)-carvone, *N*-methylmaleimide, and maleimide (Scholtissek *et al*., 2017; Tischler *et al*., 2020). Furthermore, it could be shown that FOYE-1 can reduce these compounds with the NADPH analogue 1-benzyl-1,4-dihydronicotinamide, dramatically reducing the cost for cofactors for potential future industrial applications. *Botryotinia fuckeliana* OYE isoform 4 (BfOYE4) and *Aspergillus niger* OYEs are class III fungal OYEs and could be shown to reduce cyclopentenone, cyclohexenones, and hexenal, among others (Robescu *et al*., 2022). In addition to the reduction reaction, BfOYE4 was used in an approach to isomerize α-angelica lactone (Robescu *et al*., 2022). Together with the class I OYE from *Galderia sulphuraria*, the isomerization could be achieved, leading to the (*S*)-α-angelica lactone by GsOYE and (*R*)-α-angelica lactone by BfOYE4. This is a good example of the importance of different classes in order to exploit the full potential of OYEs.

Besides the numerous studies investigating the biotechnological applications of OYEs, OPRs and their involvement in JA biosynthesis was another major focus in the field. Formation of JA starts in the chloroplast by the oxidation and cyclization of polyunsaturated fatty acids, forming the cyclopentenones, *cis*-OPDA and *dn*-*cis*-OPDA (Wasternack and Feussner, 2018). They are imported into the peroxisome, where their enone motif is reduced by an OPR and the carboxy group carrying side chain is shortened by β-oxidation, leading to the formation of JA. JA biosynthesis is triggered by different stress conditions like necrotrophic pathogens as well as feeding and wounding damage (Howe *et al*., 2018). Loss-of-function mutant lines revealed an involvement in plant fertility (Stintzi and Browse, 2000). *A. thaliana opr3* and rice (*Oryza sativa*) *Osopr7* plants are male sterile, because the pollen release from the anthers and the pollen germination are impaired (Pak *et al*., 2021; Stintzi and Browse, 2000). This phenotype can be rescued by the application of MeJA during flower development. In solanaceous plants, however, JA regulates egg-cell development and mutants are therefore female sterile (Goetz *et al*., 2012). In addition to the predominant peroxisomal OPR3 route in *A. thaliana*, a cytosolic OPR2 was found to be a secondary pathway for JA biosynthesis, explaining the residual amounts of JA in *Atopr3* plants (Chini *et al*., 2018). With a disrupted *Atopr3* gene, the carboxy group-carrying side chain of *cis*-OPDA directly undergoes three cycles of β-oxidation, yielding 4,5-ddh-JA, which can be reduced in the cytosol by OPR2 to form JA. However, this pathway is not sufficient to rescue the phenotype alone. A current study revealed that 4,5-ddh-JA could be a more mobile signal, while *cis*-OPDA is specific to the peroxisome, which could lead to a compartmentalization of JA levels and respective gene response (Mekkaoui *et al*., 2025). However, it could be demonstrated that AtOPR3 targeted to all major organelles in the *Atopr3* background could rescue the male infertile phenotype. AtOPR2 and the second cytosolic isoform AtOPR1, which prefers (-)-*cis*-OPDA over the natural (+)-*cis*-OPDA *in vitro*, could not rescue the phenotype in these organelles (Mekkaoui *et al*., 2025; Schaller *et al*., 2000; Schaller *et al*., 1998). However, when targeted to the peroxisome, both AtOPR2 and AtOPR1 could rescue the male sterile phenotype.

Here we present an extended phylogeny of the OYE superfamily throughout the domains of life and characterize an OYE from *Brevibacillus nitrificans,* which was incorrectly assigned as a *P. patens* gene in a previous genome annotation. BnOPR targeted to peroxisomes was able to rescue the male infertile phenotype of *Atopr3* plants, demonstrating the *in planta* functionality of a bacterial enzyme. The recombinant enzyme was tested for *in vitro* activity with different known OYE substrates. Applying our previously described *ex vivo* assay (Ni and Feussner, 2023), we show that BnOPR can reduce free *cis*-OPDA as well as cyclopentenone moieties esterified to membrane lipids in Arabidopsides from the *A. thaliana* metabolome. Insights into the active site of BnOPR are used to discuss potential substrate preferences compared to AtOPR2 and AtOPR3. Finally, the growth-promoting effect of *B. nitrificans*, *B. brevis, and P. physcomitrellae* on the growth of *P. patens* and the occurrence of these strains in the rhizobiome are investigated.

## Materials and Methods

### Chemicals

All chemicals used in this study were purchased from Sigma Aldrich and Carl Roth. The solvents were UHPLC-grade and purchased from Thermo Fisher Scientific and Merck Supelco. ddH2O water was obtained from a Satorius Millipore device. If not stated otherwise, the Kits and restriction enzymes used for molecular cloning were obtained from Marcherey-Nagel, Thermo Fisher Scientific, or New England Biolabs.

### Multiple sequence alignment and structure prediction

Amino acid sequences for BnOPR (WP_328204078, annotated as Pp3s58_150 before contamination removal (Lang *et al*., 2018)), AtOPR1 (At1g76680), AtOPR2 (At1g76690), AtOPR3 (At2g06050), Rer-ER7 (QBR53095.1), Chr-OYE1 (ALE60336.1), and Ppo-ER3 (WP_330721231.1) were subjected to multiple sequence alignment analysis using Clustal Omega (Madeira *et al*., 2024). Structures for BnOPR and AtOPR2 were predicted using Alphafold2 Colabfold (Mirdita *et al*., 2022). Visualization of the structural predictions, as well as AtOPR1 (PDB: 1VJI) (Fox *et al*., 2005) and AtOPR3 (PDB: 2G5W) (Han *et al*., 2011) was performed with Pymol (Schrödinger Inc.).

### Maximum-likelihood phylogeny

For phylogenetic studies, AtOPR3 protein sequence was used as a query to screen against the protein data 438 genomes, consisting of 38 plant genomes (For a detailed list: Supplementary information), the top 100 hits against Paenibacillaceae, the top 100 hits against non-Paenibacillaceae, the top 100 hits against non-Paenibacillaceae and non-Bacillaceae, and the top 100 hits against non-Bacteria. The hits were selected, aligned using MAFFT L-INS-I (Katoh and Standley, 2013), and a maximum-likelihood phylogeny was computed with 100 bootstrap repeats, using IQ-TREE multicore version 1.5.5 (Nguyen *et al*., 2015); using Modelfinder (Kalyaanamoorthy *et al*., 2017), 144 protein models were tested and the best-fit model was LG+G4, chosen according to Bayesian Information Criterion. The tree was visualized in iTOL (Letunic and Bork, 2024) and colored according to the existing clades.

### Protein expression and purification

BnOPR CDS was inserted into pET28 (+) from BioCat GmbH with the restriction sites NdeI and BamHI. The alignment included a TetR7AcR family transcriptional regulator at the N-terminus, which was considered as an error in the annotation. The truncated version, containing the BnOPR domain, was amplified, digested with NdeI and BamHI, and ligated into a respective digested pET28 (Merck, Darmstadt, Germany) backbone. The full-length construct was transformed into *E. coli* BL21 (DE3). Expression and purification were conducted as described (Mohnike *et al*., 2021). Protein was further purified by size exclusion chromatography (SEC) with a Sephadex S200 26/60 column (Cytiva, Chicago, IL, USA) equilibrated in 50 mM HEPES (pH 7.5), 50 mM NaCl, 10 % (*v/v*) glycerol, and 2 mM MgCl2. Protein was concentrated to approx. 10 mg/mL using a Centricon Spin column (MWCO 10 kDa). AtOPR3 and AtOPR2 CDS inserted into pET28 (+) were ordered from NovoGene with the restriction sites EcoRI and NotI as well as NheI and HindIII, respectively. AtOPR1/2/3 purification was described previously (Mekkaoui *et al*., 2025). For AtOPR1, buffer was exchanged by SEC with a Sephadex S200 26/60 column equilibrated in 50 mM Tris/HCl (pH 7.8), 50 mM NaCl, 10 % (*v/v*) glycerol, and 2 mM MgCl2. For AtOPR2 and AtOPR3 buffers were exchanged using a HiPrep 26/60 desalting column equilibrated in 50 mM Tris/HCl (pH 7.8), 50 mM NaCl, 10 % (*v/v*) glycerol, and 2 mM MgCl2. Enzymes were used directly for kinetic measurements or flash frozen in liquid nitrogen and stored at −80 °C until further use in *in vitro* or *ex vivo* screens.

### *In vitro* assays and kinetics

For kinetic measurements, enzymes were used directly after the purification. Enzyme concentration was measured, using the FMN extinction of denatured enzyme in 6 M Guanidin-HCl and a respective molar extinction coefficient of 12,913 M^-1^cm^-1^ at 446 nm. Purified enzyme was frozen in liquid nitrogen and stored at −80 °C until usage. Frozen enzyme aliquots were thawed on ice immediately before the assay. 0.1 mg/mL BnOPR or AtOPR3 were mixed carefully with 10 mM NaCl, 1 mM NADPH, and 1 mM MgCl2 in 20 mM Tris/HCl (pH 7.8). After the reaction mixtures reached 30 °C, the reactions were started by the addition of 0.1 mM *cis*-OPDA. Negative controls were conducted with heat-inactivated enzyme. Reactions were either stopped after 10 min or 90 min by the addition of 1x reaction volume acetonitrile. Precipitated enzymes were removed by centrifugation at 18.000 x g for 15 min before analysis by ultra-high-pressure liquid chromatography coupled to high-resolution mass spectrometry (UHPLC-HRMS, Agilent Technologies Inc.) (Feussner *et al*., 2023). Further assays were set up accordingly, using the following substrates (with respective concentrations): 4,5-ddh-JA (50 μM), prostaglandin A2 (100 μM), cortisone (300 μM), prednisone (300 μM), menadione (0.5 mM), 2,4-dinitrophenol (2,4-DNP)(0.5 mM), 2-cyclohexen-1-one (1 mM), duroquinone (1 mM), maleic acid (1 mM), 3-hydroxy-4,6-dimethyl-cyclohex-2-enone (1 mM), methylvinylketone (1.5 mM), citral (3 mM), 2-cyclopenten-1-one (3 mM), (2*E*,4*E*)-hexadienal (3 mM), (2*E*)-hexenal (3 mM), 2-methyl-2-cyclopentenone (3 mM), (2*E*)-pentenal (3 mM), cinnamaldehyde (3 mM), and *cis*-OPDA (50 μM). For kinetic measurements, 0.25 mM NADPH, 12.5 U/mL glucose oxidase, 20 mM glucose, and 10 mM NaCl were mixed in 20 mM HEPES (pH 7.5) with 0-10 mM substrate. 1-4 µM freshly purified enzyme was used to start the reaction. Absorbance change at 340 nm and 25 °C was observed with a Thermo Genesys 50 spectrophotometer (Thermo Fisher Scientific Inc.). Data was plotted using Origin (OriginLab Corporation) and fitted accordingly to receive kinetic parameters.

### Metabolite extraction and *ex vivo* assay

*Atopr2 Atopr3* plants were sprayed with 500 μM MeJA, wounded 0.5 h later by squeezing each leaf three times with forceps and harvested after one hour to be directly frozen in liquid nitrogen. Samples were stored at −80 °C until usage. Plant material was ground and 150 mg were weighed in per sample while still being frozen. Metabolites were extracted using an MTBE two-phase extraction as described in (Feussner *et al*., 2023). The dried metabolites were dissolved in 10 mM Tris/HCl (pH 7.8), adding 1 mM NADPH, 4 mM NaCl, and 1 mM MgCl2. Reactions were strated b the addition of 4 μM enzyme and were incubated for 2 h at 30 °C. The reactions were stopped by adding one volume methanol, samples were centrifuged at 18.000 x g for 15 min at 4 °C, and transferred into LC-MS vials. Samples were analyzed UHPLC-HRMS. Peak picking and peak alignment were conducted with Profinder (Agilent Technologies Inc.), and statistical calculations and clustering were performed using the MarVis Suite toolbox (Kaever *et al*., 2009)

### *Atopr3* complementation and analysis

For stable expression of BnOPR in *Atopr3* plants, a plant transformation vector was generated. Full-length BnOPR CDS, including the N-terminal transcriptional regulator (TetR7AcR family transcriptional regulator) and a peroxisomal targeting sequence (SKL) at the C-terminus, was cloned into a pENTRY-E vector carrying a *CaMV35S* promoter. Using Gateway technology (Thermo Fisher Scientific) as described previously (Heilmann *et al*., 2012), the binary vector *pCAMBIA33-CaMV35S::BnOPR-SKL* was created, which is carrying the resistance gene against glufosinate. Transgenic plants were generated through *Agrobacterium tumefaciens*-mediated transformation via floral dipping (Clough and Bent, 1998). As *Atopr3* plants are male sterile, the plants were daily sprayed with 500 µM MeJA as soon as flower buds were appearing until 3 weeks after transformation. Selection of transgenic T1 plants was performed by herbicide treatment (Basta, Bayer CropScience). Silique formation was monitored and seeds from plants with restored fertility were collected for further experiments.

Complementing lines, wild-type (WT) cv. WS and *Atopr3* plants were grown under long-day conditions for four or six weeks. After four weeks, plants were wounded by squeezing every leaf three times with a tweezer. After one hour, leaves were collected and shock frozen in liquid nitrogen. Until extraction, samples were stored at −80 °C. Plant material was ground into a fine powder, while still being frozen, weighed into 100 mg, extracted and analyzed by ultra-high-pressure liquid chromatography coupled to tandem mass spectrometry (UPLC-MS/MS) as described (Herrfurth and Feussner, 2020).

### *P. patens* growth experiments

*P. patens* ecotype Gransden 2004 was grown on solid BCD-AT under long-day conditions with a light intensity of 70 µmol/m^2^s at 22 °C. *B. nitrificans* was acquired from DSMZ (DSM No.: 26674) and was grown on nutrient agar. Overnight cultures in liquid nutrient medium were grown at 37°C and 200 rpm to be washed with sterile ddH2O and diluted to 25,000 cells/mL. 20 µL were used to inoculate *P. patens* protonema spots on solid minimal medium supplemented with 5 % (*w/v*) sucrose and 1.5 % (*w/v*) plant agar (Bécard and Fortin, 1988). Plants were imaged and measured after 25 days. *P. patens* ecotype Reute was grown on solid Knop medium in petri dishes under long-day conditions with a light intensity of 70 mol/m^2^s at 22 °C (Egener *et al*., 2002). *B. brevis* was acquired from JCM (JCM no.: 6285), cultivated on LB-plates at 37 °C, and liquid LB cultures were inoculated with single colonies and grown at 37 °C overnight at 200 rpm. *P. physcomitrellae* was acquired from DSMZ (DSM no.: 29851) and was grown at 28 °C on TSB medium. Cell density was determined and diluted to 25,000 cells/mL with ddH2O. *B. brevis, P. physcomitrellae,* and *P. patens* were co-cultivated on solid minimal medium supplemented with 5 g/L sucrose and 2 g/L phytagel. For gametophore growth experiments, *P. patens* colonies were inoculated with 20 µL (500 cells) of either *B. brevis* culture or *P. physcomitrellae*. For rhizoid growth experiments, mature gametophore tops were picked and transferred to new plates before inoculation. Petri dishes were closed with Parafilm and incubated for 25 days under long-day conditions.

## Results

### BnOPR clusters within the distinct bacterial clade of OYE

In order to identify OYEs in bryophytes, a phylogenetic analysis of the OYE enzyme family throughout the domains of life was carried out. This analysis made use of the extensive sampling of streptophyte genomes (Kunz *et al*., 2025), recovering a fully supported clade of streptophyte sequences with the land plants nested within. Land plant OYE homologs are distributed between two deeply split major clades (Figure 1). To our surprise, *P. patens* gene, Pp3s58_150 (genome V3) fell into the bacterial clade of OYEs (Figure 1, BnOPR)(Lang *et al*., 2018). A BLAST analysis, however, revealed the existence of this gene in the genome of *B. nitrificans* with a length of 1761 bp. It included not only the OYE domain but a transcriptional regulator from the TetR7AcR family (Figure S1A)(Tatusova *et al*., 2013). First *in vitro* activity tests with the entire recombinant protein confirmed that the enzyme activity was not impaired by the transcriptional regulator. A closer analysis of this gene, however, revealed a second start codon leading to a truncated version of 1107 bp. This protein was called BnOPR and analyzed by kinetics and *ex vivo* assays. Along this line, this gene was recently removed from version 6 of the *P. patens* genome (Bi *et al*., 2024). Based on the occurrence of *BnOPR* within earlier assemblies of the *P. patens* genome, an co-isolation of both species is assumed. This reneders the possibility of *B. nitrificans* to be plant-associated likely.

**Figure 1.**
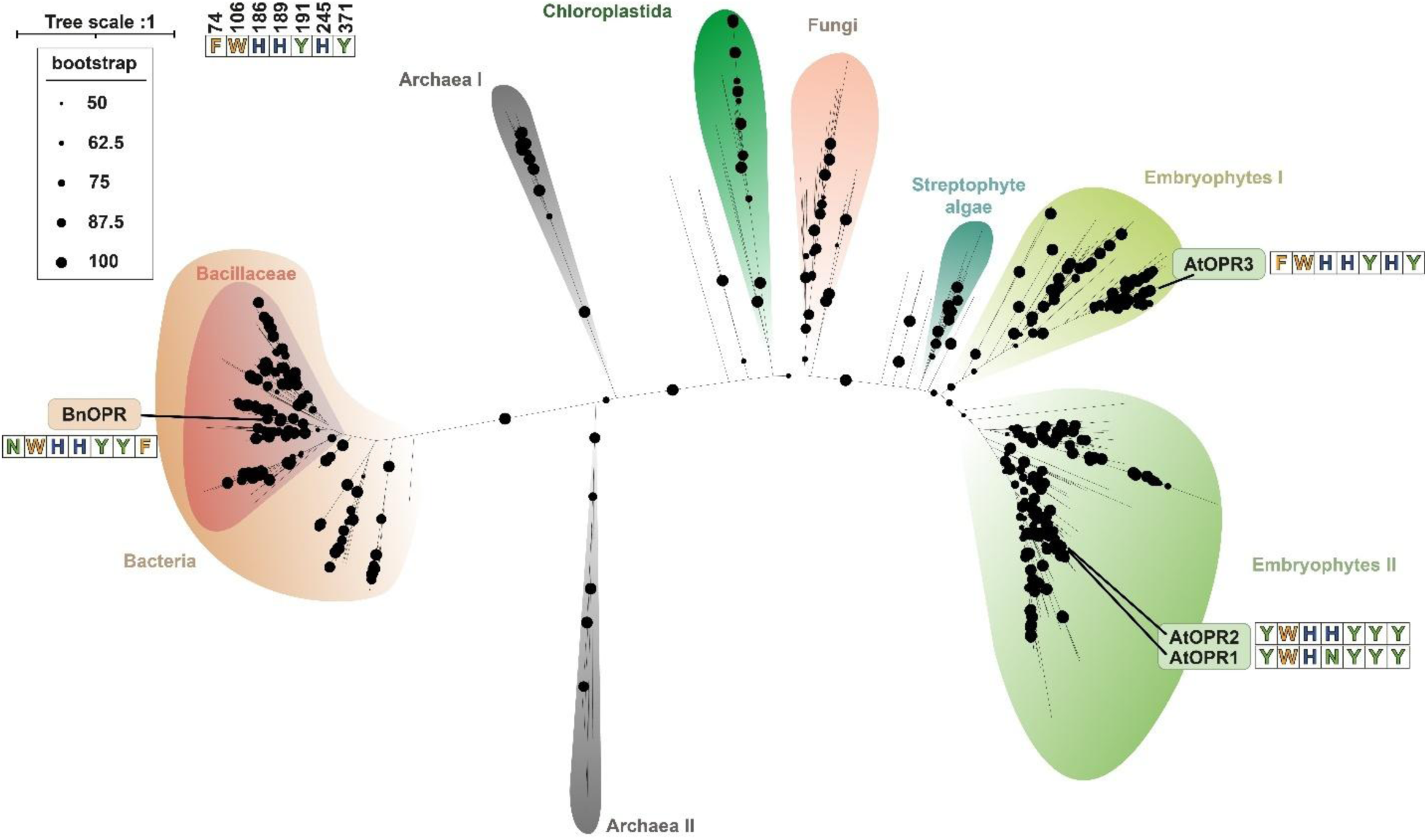
Phylogenetic tree of OYEs from 438 organisms. Unrooted phylogenetic tree of OYEs in 438 plant, bacterial, and archaeal genomes. Black circles indicate the bootstrap support of each branch with the respective size as stated in the legend. AtOPR1-3 and BnOPR are marked in their respective clades with important active site residues marked according to their polarity (basic: blue, polar: green, unpolar: orange). The number of the residue, as marked in the legend, is according to AtOPR3. **Alt text:** Phylogenetic tree of Old Yellow Enzyme protein sequences throughout the domains of life. The distinct clades were labelled according to the origin domain in the tree of life. The clades from left to right are Bacteria, Archaea I, Archaea II, Chloroplastida, Fungi, Streptophyte algae, Embryophytes I, and Embryophytes II. The bacterial clade has a subdivision of Bacillaceae. The candidates *Brevibacillus nitrificans* BnOPR, AtOPR1, 2, and 3 are highlighted in the clades Bacteria subclade Bacillaceae and Embryophytes I and II, respectively. For comparison, the active site residues, according to Breitahupt et al. (2009), are mentioned for each candidate.

The amino acid sequence of BnOPR shows a 30.28 % similarity to yqjM from *Bacillus subtilis* in the same clade (Fitzpatrick *et al*., 2003). AtOPR1, AtOPR2, and AtOPR3 have 33.14 %, 33.43 %, and 28.41 % similarity, respectively (Schaller *et al*., 2000; Schaller *et al*., 1998) (Figure S1C). 54 % to 65 % similarity was discovered with class IV OYEs from *Crysoebacterium* sp. CA49, *Pyricularia oryzae,* and *Rhodococcus erythropolis* (Peters *et al*., 2019). BnOPR belongs to the class IV OYEs based on the residues interacting with the FMN cofactor and the 18 amino acid loop extension between α-helix 7 and β-sheet 7. Apart from this extension, class IV and class II OYEs differ mainly by the C-terminal residues for monomer-monomer interaction, which are absent in BnOPR (Adalbjörnsson *et al*., 2010; Peters *et al*., 2019; Shi *et al*., 2020). The active site residues, which were described to be important for substrate binding and catalysis, were further investigated by multiple sequence alignment, based on the results for OPRs (Breithaupt *et al*., 2009). Seven out of nine residues in the FMN binding site are conserved between AtOPRs and BnOPR (Figure S1B). The latter has a Gly321Lys and a Phe370Pro exchange. Tyr191, which is important for catalysis, is conserved as well. Residues for substrate binding and substrate specificity show some differences. For AtOPRs, substrate specificity is mediated by Tyr74 and Tyr242 in AtOPR1 or Tyr76 and Tyr244 in AtOPR2, and Phe74 and His245 in AtOPR3. BLAST results indicated Asn72 and Tyr233 for BnOPR, for which the positioning and orientation within the model suggests otherwise (Figure 2). Tyr233 faces towards the active site, but shows some increased distance to the active site compared to the AtOPRs. Asn72 on the other hand faces outwards of the active site. The substrate binding site is conserved in four out of five residues. The fifth residue Tyr371 is replaced by Val368 in BnOPR, according to the sequence alignment. However, Val368 is the C-terminal residue of BnOPR and is not oriented towards the active site, according to the predicted structure. Respectively, the substrate binding site shows some conservation, but the residues mediating substrate specificity towards *cis*-OPDA are exchanged. This is also represented in the FMN binding site and surface models (Figures S2+S3). While AtOPR3 has a pocket presenting the FMN cofactor, AtOPR2 has a narrower binding pocket covering most of the FMN. The BnOPR model is more comparable to AtOPR3, however, the binding site is even more exposed to the surface. The increased exposure of FMN is a result of the outwards facing loop, containing Phe371. Based on the phylogenetic distance of BnOPR to the AtOPRs, as well as the different active site geometry, amino acids involved and overall spacing, it is unlikely that the occurrence of BnOPR in *B. nitrificans* is a result of a horizontal gene transfer event.

**Figure 2.**
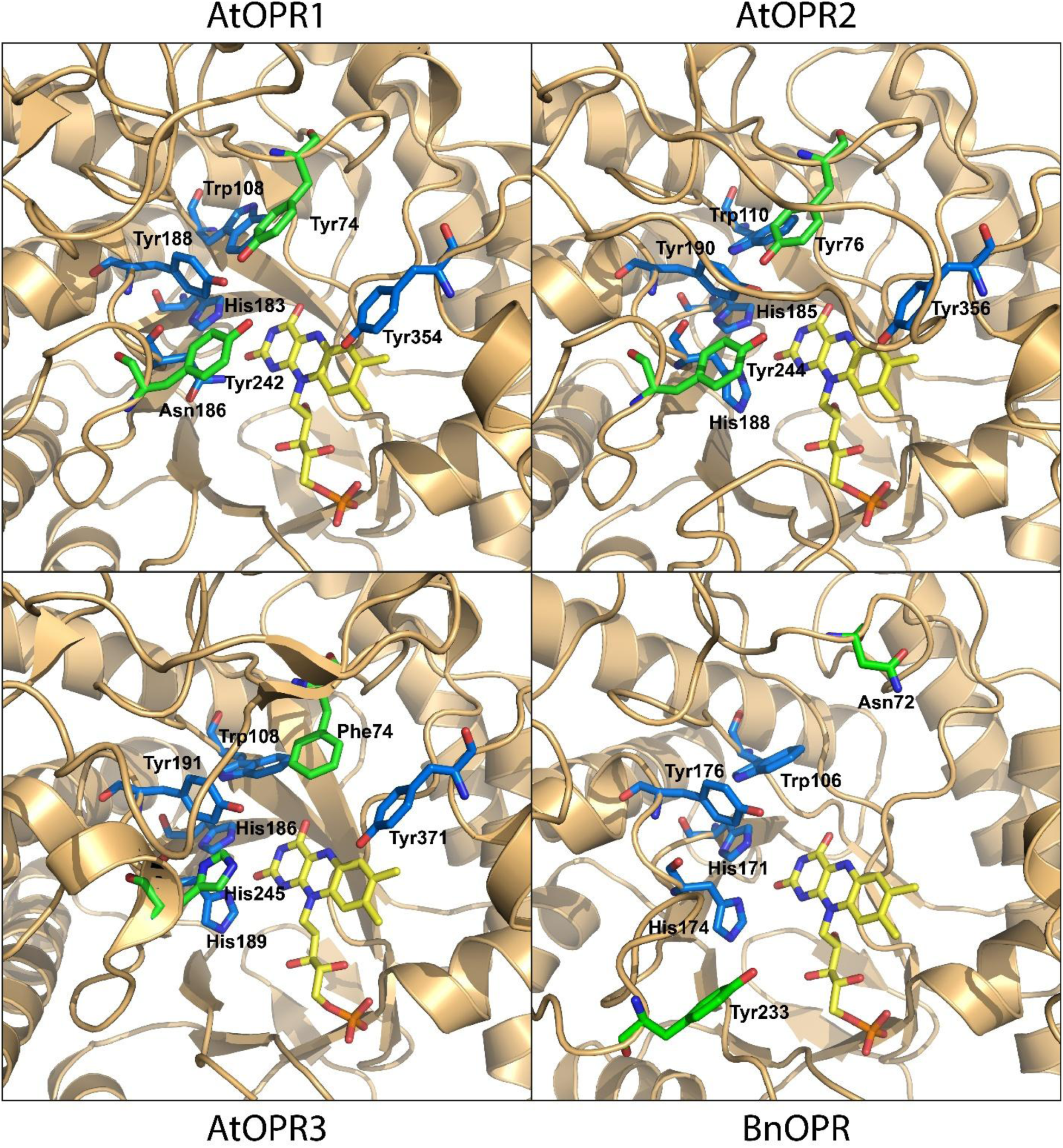
Comparison of the active sites of BnOPR and AtOPR1-3. Comparison of active site residues from the crystal structures of AtOPR1 (PDB: 1VJI) and AtOPR3 (PDB: 2G5W). BnOPR and AtOPR2 were modeled using Alphafold2 Colabfold (Mirdita *et al*., 2022). Residues for substrate binding and catalysis are colored blue, and residues that determine substrate specificity are colored green. Residues and numbers refer to the position in the respective protein. The FMN cofactor is displaed in yellow sticks. **Alt text:** Displayed are four panels with the active site residues for substrate binding and catalysis for AtOPR1, 2 and 3 as well as BnOPR. The enzyme backbone is shown in beige, and the FMN cofactor is displayed in yellow. The structures are based on X-ray crystal structures for AtOPR1 and AtOPR3 or modelled by Alphafold for AtOPR2 and BnOPR. The two residues determining substrate specificity for the OPDA enantiomers are shown in green and differ between the enzymes. While AtOPR1 and AtOPR2 have two Tyr residues, AtOPR3 has a Phe and a His. For BnOPR, a Tyr and an Asn are shown at the respective positions. Further residues for substrate binding and catalysis are shown in blue sticks. Heteroatoms nitrogen are shown in dark blue, oxygen in red, and phosphate in orange.

### BnOPR complements *Atopr3* male infertility phenotype

BnOPR coding sequence with a length of 1761 bp was fused C-terminally with a SKL-peptide sequence for peroxisomal import and used in complementation experiments with *Atopr3*. Mutant plants were transformed with pCambia vectors containing the TetR7AcR-BnOPR CDS with a *CaMV35S* promoter by floral dipping via *A. tumefaciens*. Transformed plants were checked for flower development and silique formation. As described previously, *Atopr3* plants are male sterile, with shorter filaments and anthers still intact. Complementing lines with *CaMV35S:: TetR7AcR-BnOPR-SKL,* produced siliques again and showed the wild-type phenotype (Figure 3). To confirm that the plants produce more JA-Ile compared to the *Atopr3* plants, metabolites were extracted from wounded leaves and analyzed by quantitative UPLC-MS/MS (Figure 4). JA levels are decreased by 95 % in the *Atopr3*. Compared to the mutant, the complementations show a 5-fold increase in JA and JA-Ile levels with slight variations between the lines and correspond to about 20 % of the wild-type levels. Hydroxy-JA and hydroxy-JA-Ile, however, reach up to 75 % of WT levels. These wound-induced levels of JA and JA derivatives in the complemented *Atopr3* lines, together with the WT flower phenotype, demonstrated the OPR activity of TetR7AcR-BnOPR *in planta*.

**Figure 3.**
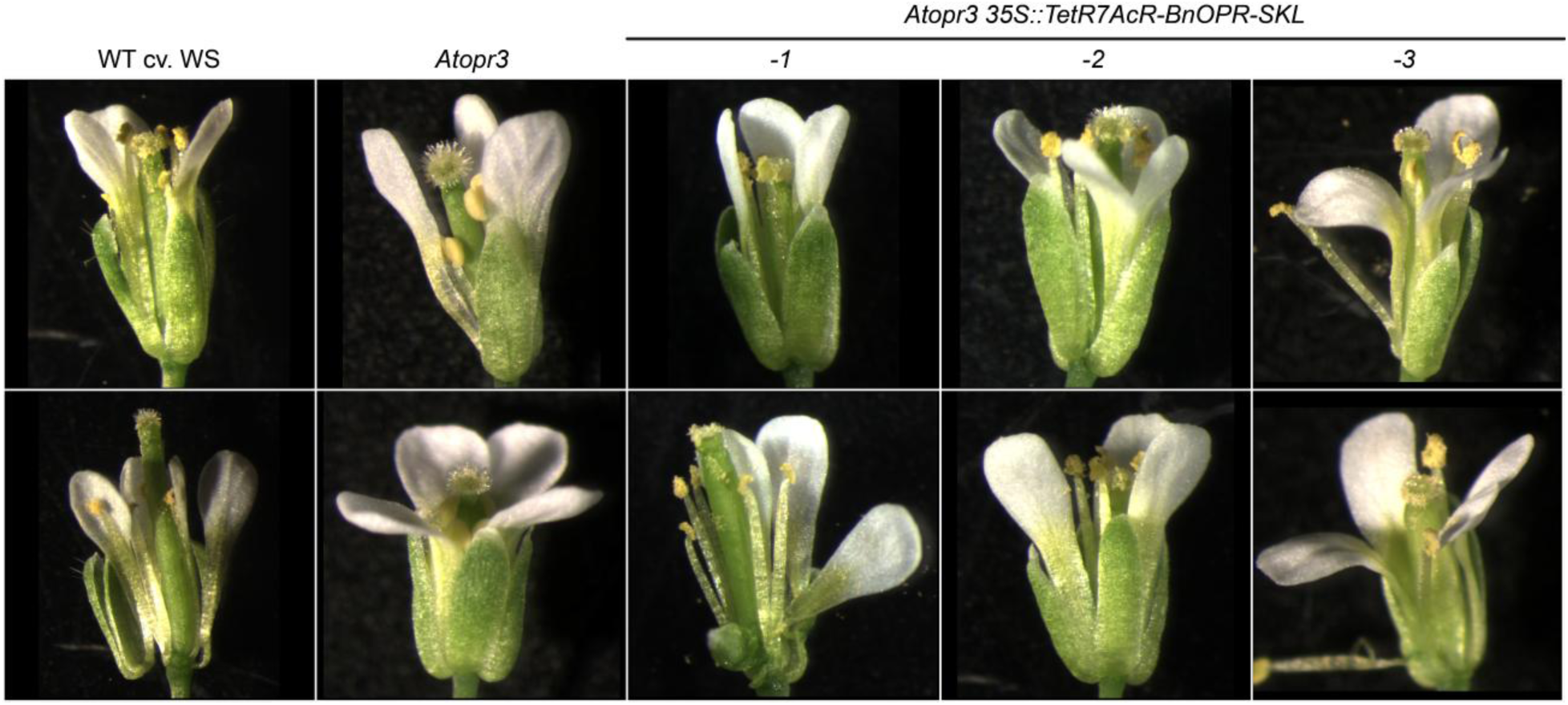
Flowers of *Atopr3* and complementing lines with BnOPR. *Atopr3* mutant lines were complemented by floral dipping using *A. tumefacins* transformed with *pCAMBIA33-CaMV35S::TetR7AcR-BnOPR-SKL*. Transformants were selected via herbicide resistance. All lines are in the ecotype Wassilewskija. For each line, three biological replicates were grown and analyzed. **Alt text:** The ten panels show *Arabidopsis thaliana* flowers on black background. The wild-type flowers on the left, in the ecotype Wassilewskija, show normal physiology. The anther filament elongation is intact and the pollens are released to fertilize the stigma. The two panels to the right of the wild type show male infertile *Atopr3* mutant line flowers. There, the anther filament elongation is impaired, and the anthers appear smooth because the pollen is not released. The six panels to the right show three independent lines of *Atopr3*, which were transformed to transiently express *BnOPR* with the peroxisomal targeting signal SKL under the control of a *CaMV35S* promoter. Here, the anther filament elongation is restored, and flowers appear wild-type-like again.

**Figure 4.**
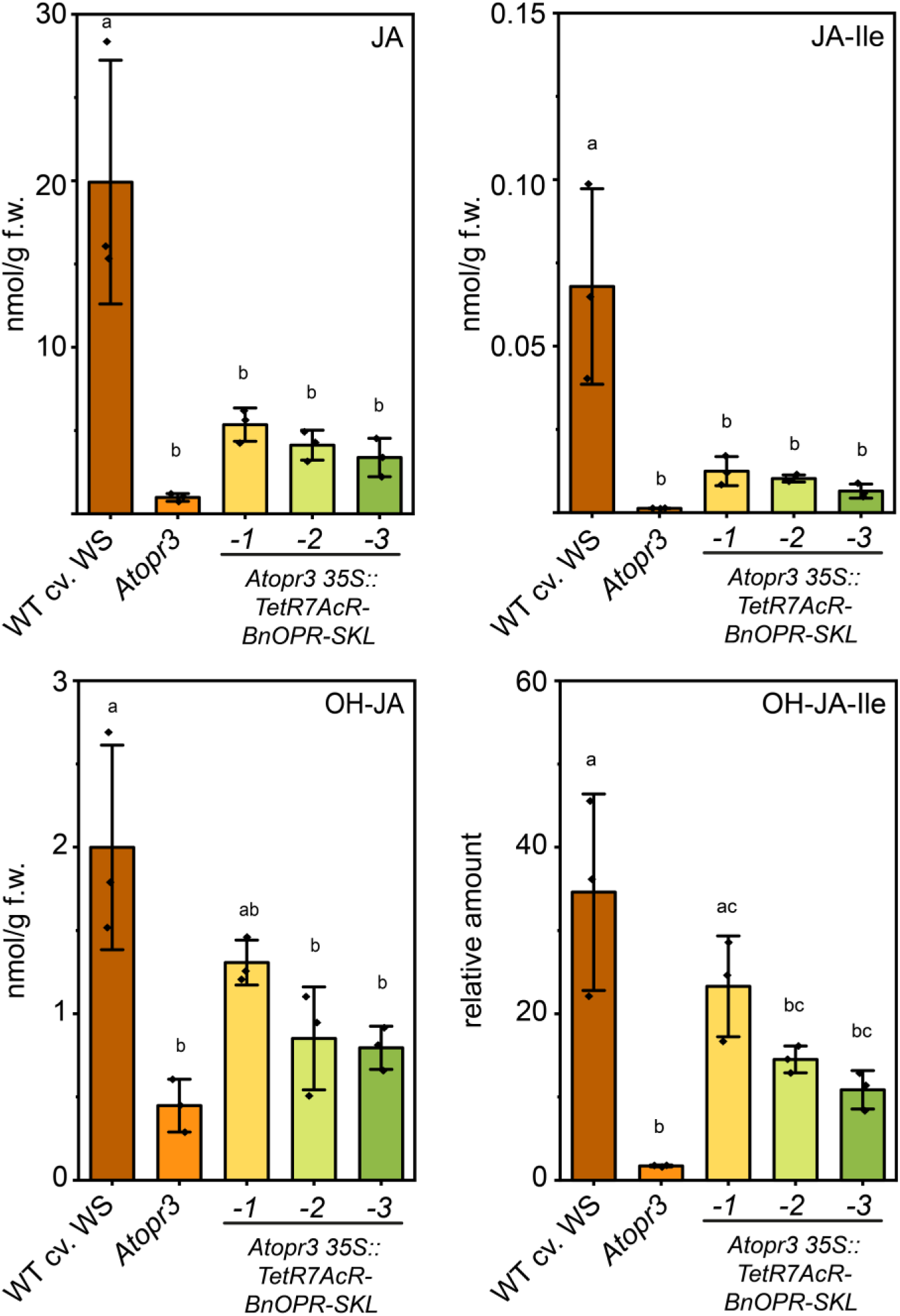
Phytohormone profiling in complemented *Atopr3* lines. Absolute amounts of JA, JA-Ile, hydroxy-JA, and hydroxy-JA-Ile after two-phase extraction of leaf material, analyzed by quantitative UPLC-MS/MS. Leaves were wounded and harvested 1 hour post wounding. Plant material was flash frozen in liquid nitrogen, ground, weighed, and extracted. Polar and unipolar phases were pooled after extraction, evaporated, and dissolved in MeOH:H2O (4:1). Samples were analyzed with UPLC-MS/MS as described (Herrfurth and Feussner, 2020). Data were acquired from three biological replicates. Statistics for three biological replicates were calculated by one-way ANOVA followed by *post-hoc* Tukey’s HSD test and are indicated by different letters (*P*<0.05). **Alt text:** The four bar graphs show phytohormone levels in WT, *Atopr3*, and complementing lines. Shown are jasmonic acid (top left), jasmonoyl-isoleucine (top right), hydroxy-jasmonic acid (bottom left) and hydroxy-jasmonoyl-isoleucine (bottom right). All phytohormone levels are strongly reduced in *Atopr3*. The complementing lines show increased levels compared to the mutant lines, showcasing the succesful complementation.

### BnOPR reduces a variety of α,β unsaturated enones but no nitro compounds

To characterize the activity of BnOPR *in vitro*, the enzyme was heterologously expressed in *E. coli* and purified by affinity chromatography and SEC. To investigate the buffer conditions best suited for kinetics, a thermal shift assay was conducted (data not shown). Highest stability was achieved in HEPES pH 7.5, which was subsequently used for *in vitro* reactions and kinetics. The purified enzyme was tested with *cis*-OPDA, prostaglandin A2, 4,5-ddh-JA and 3,7-ddh-JA as different cyclopentenone substrates (Figure 5). Prednisone and cortisone were tested to assess the reduction of steroles and positional preference of BnOPR. Finally 2,4-DNP was tested to investigate nitro group reduction. After ten or 90 min, respectively, the reactions were stopped and analyzed by UHPLC-HRMS. Conversion was evaluated based on substrate consumption and product formation independently of their different ionization efficiencies.

**Figure 5.**
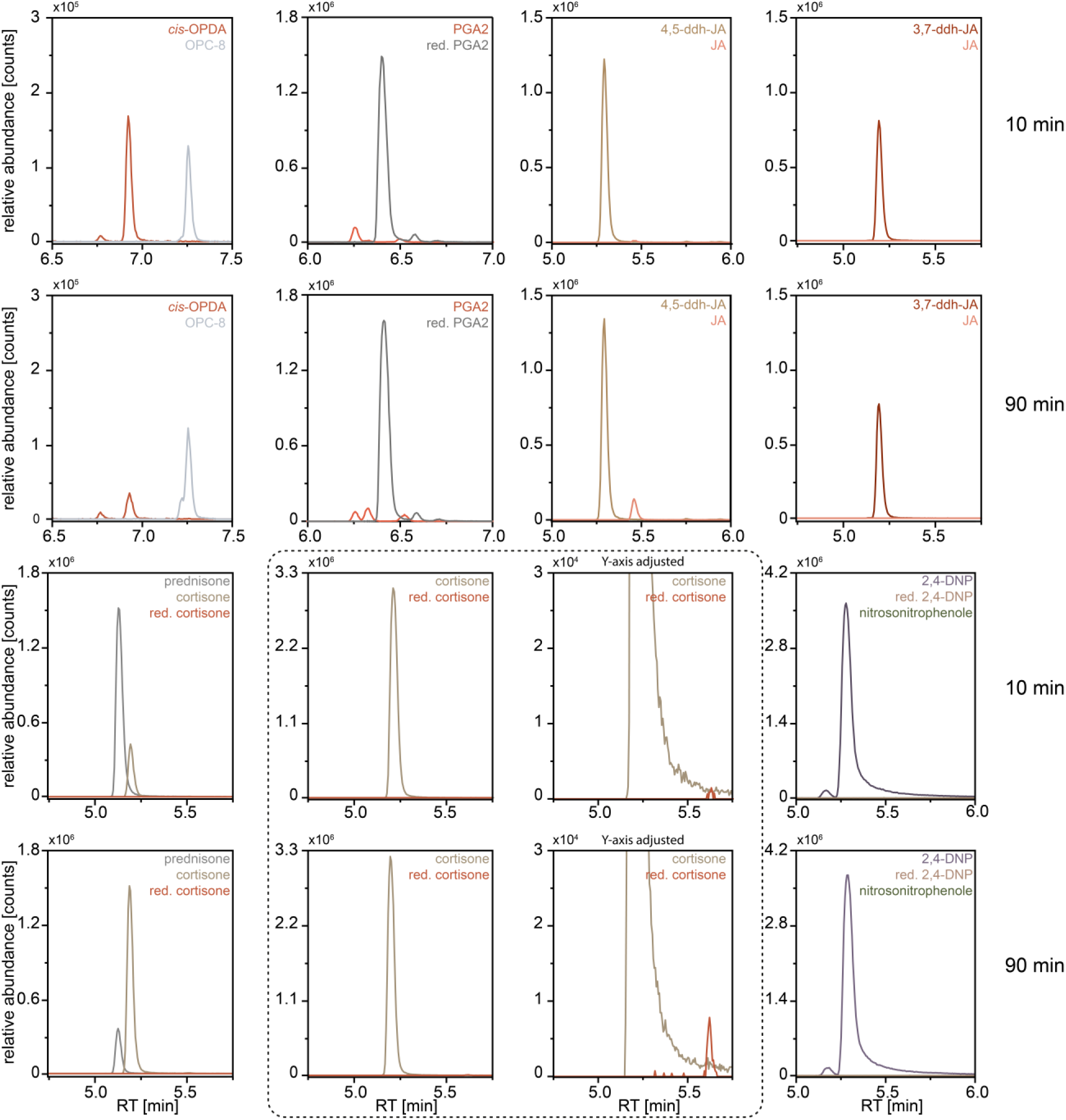
BnOPR reduces *cis*-OPDA *in vitro. cis*-OPDA, prostaglandin A2 (PGA2), 4,5-ddh-JA, 3,7-ddh-JA, prednisone, cortisone and 2,4-dinitrophenole (2,4-DNP) were incubated with heterologously expressed and purified BnOPR. Reactions were incubated at 30 °C and stopped after 10 and 90 min by the addition of one volume acetonitrile. After centrifugation to remove precipitated protein, assays were measured by UHPLC-HRMS and analyzed towards substrate consumption and product formation. Graphs show the extracted ion chromatogram of all substrates with their respective products. Respective extracted *m/z* values: [*cis*-OPDA+H^+^]^+^ 293.211, [OPC-8+H^+^]^+^ 295.227, [PGA2-H^+^]^-^ 333.209, [red. PGA2-H^+^]^-^ 335.224, [4,5-ddh-JA+H^+^]^+^ 209.117, [JA+H^+^]^+^ 211.133, [3,7-ddh-JA+H^+^]^+^ 209.117, [prednisone+H^+^]^+^ 359.186, [cortisone+H^+^]^+^ 361.201, [red. cortisone+H^+^]^+^ 363.217, [2,4-DNP-H^+^]^-^ 183.006, [put. red. 2,4-DNP-H^+^]^-^ 185.020, [put. nitrosonitrophenol-H^+^]^-^ 167.010. Figures are representative of three measurements. **Alt text:** The graphs show extracted ion chromatograms of different Old Yellow Enzyme substrates, which were incubated with purified enzyme to test for conversion. The reactions were analyzed by UHPLC-HRMS, and expected mass per charge signals were extracted. The top row shows the reactions, from left to right, for *cis*-OPDA, prostaglandin A2, 4,5-didehydro-jasmonic acid, and 3,7-didehydro-jasmonic acid after ten min. The second row shows the same reactions after 90 min. Here, the conversion of *cis*-OPDA, prostaglandin A2, and to a lesser extent 4,5-didehydro-jasmonic acid could be shown. 3,7-didehydro-jasmonic acid is no substrate of BnOPR. The third and fourth rows show the reactions with prednisone and cortisone, a close-up of the cortisone reaction, and 2,4-DNP after 10 or 90 min. Here, prednisone acts as substrate, while cortisone showed just minor conversion, and 2,4-DNP was not converted.

In case of *cis*-OPDA and 4,5-ddh-JA, conversion was compared to the activity of AtOPR3 and AtOPR2. AtOPR3 fully converted 0.1 mM *cis*-OPDA, while using up only around a third of the 4,5-ddh-JA. AtOPR2 shows an inverse trend: it fully reduced 4,5-ddh-JA to JA but only around 2-3 % of the *cis*-OPDA (Mekkaoui *et al*., 2025). By comparison BnOPR used up 80 % of *cis*-OPDA and 15 % of 4,5-ddh-JA, placing it in between AtOPR3 and AtOPR2 activity. Therefore, additional putative substrates were tested (Table S1). AtOPR3 and AtOPR2 reduced 2-cyclohexen-1-one, cinnamaldehyde, and citral to a similar extent. 2-cyclopenten-1-one, maleic acid and (2*E*,4*E*)-hexadienal were only reduced by AtOPR3. AtOPR3 reduced with higher efficiency than BnOPR menadione, pentenal, and hexenal. On the other hand, BnOPR reduced 3-methyl-2-cyclopenten-1-one and prednisone, while AtOPR3 exhibited no activity towards these substrates. To test the ability for full conversion by BnOPR, further substrates were tested for 10 and 90 min (Figure 5). Prostaglandin A2 was fully converted after 10 min. 3,7-ddh-JA was tested but could not be reduced by BnOPR. Further, to test which double bond is preferably reduced in prednisone, cortisone was used as a substrate and showed just minor reduction, revealing the positional preference of BnOPR.

To measure exact conversion efficiencies for BnOPR, continuous kinetics were recorded with selected substrates by monitoring the NADPH consumption at 340 nm. Absorbance change was plotted against the used substrate concentration and fitted according to Michaelis-Menten equation. The structures of the substances tested, as well as the measured constants, are shown in Table 1. Turnover numbers kcat were calculated using the fitted Vmax and enzyme concentration. The KM of BnOPR for *cis*-OPDA was 53 ± 12 µM with a turnover number of 0.59 ± 0.02 s^-1^. Prednisone was bound with a KM of 232 ± 41 µM but was metabolized faster, as shown by a turnover of 1.89 ± 0.08 s^-1^. In theory, prednisone has two double bonds which can be reduced by OYEs. However, a reduced product, which lacked both double bonds, could not be detected via UHPLC-HRMS analysis. KM values for other substrates were in the mM range but showed higher turnover numbers. A preference for cyclic or aliphatic enones could not be observed. 2,4-DNP, 3,5-dinitrosalicylic acid, and 3,7-ddh-JA were tested but not reduced.

**Table 1.**
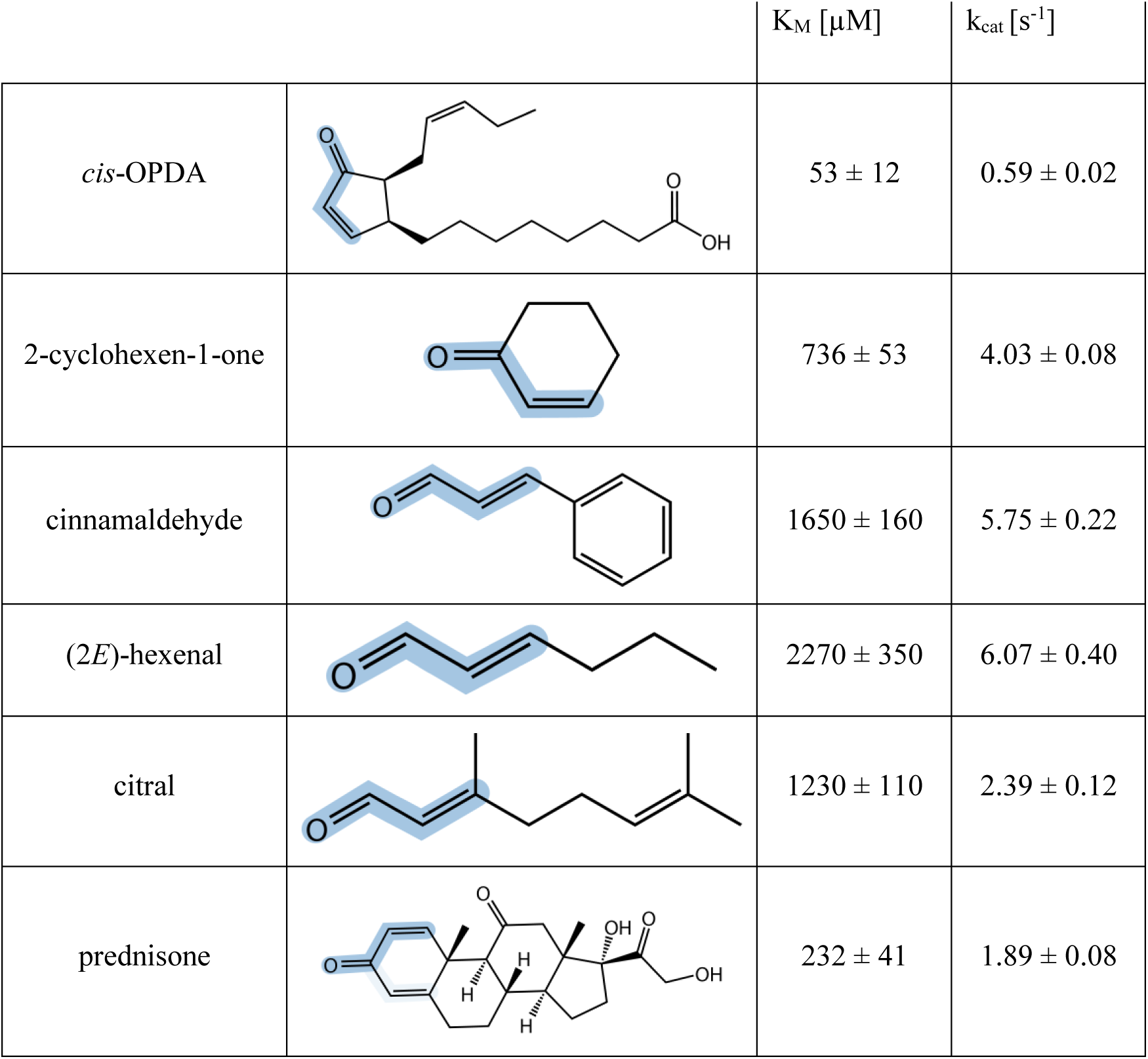
Structures of several α,β-unsaturated enones and kinetic parameters with BnOPR. Heterologously expressed and purified BnOPR was used in kinetic measurements at pH 7.5 and 25 °C. Absorbance change at 340 nm was observed using a Thermo Genesys 50, plotted and fitted with Origin. Reduced double bonds are highlighted in blue in the structures (left). Kinetic parameters are shown right hand side and were calculated based on three independent measurements.

| | | $K_M$ [μM] | $k_{cat}$ [$s^{-1}$] |
| --- | --- | --- | --- |
| *cis*-OPDA | | $53 \pm 12$ | $0.59 \pm 0.02$ |
| 2-cyclohexen-1-one | | $736 \pm 53$ | $4.03 \pm 0.08$ |
| cinnamaldehyde | | $1650 \pm 160$ | $5.75 \pm 0.22$ |
| (2*E*)-hexenal | | $2270 \pm 350$ | $6.07 \pm 0.40$ |
| citral | | $1230 \pm 110$ | $2.39 \pm 0.12$ |
| prednisone | | $232 \pm 41$ | $1.89 \pm 0.08$ |

### BnOPR reduces Arabidopsides among others in *Atopr3* metabolite extract

To identify additional substrates of BnOPR in plants, extracts from MeJA treated and wounded leaves of the *Atopr2 Atopr3* mutant, which should accumulate preferentially OPR substrates, were incubated either with recombinant AtOPR1, AtOPR2, AtOPR3, or BnOPR. Extracts were analyzed via UHPLC-HRMS, and the resulting one-dimensional self-organizing maps (1D-SOMs) revealed several clusters representing a substrate or product pattern for BnOPR activity (Figure 6). Comparing the overall cluster pattern, BnOPR was more similar in respect to substrate consumption and product formation to AtOPR3 than to AtOPR1 or AtOPR2. Analyzing the conversion of OPDA as a positive control revealed that BnOPR converted it to a similar extent as AtOPR3, while AtOPR1 and AtOPR2 showed residual amounts of OPDA. 4,5-ddh-JA, as an *in vitro* and *in planta* confirmed substrate of AtOPR2, was also reduced by AtOPR3, but neither by BnOPR nor AtOPR1. The chemical structures of additional features of interest were investigated by MS/MS fragmentation. Among them were Arabidopsides A and B, a flavonoid, as well as hexosyl glycerol containing OPDA or dn-OPDA derivatives could be identified as substrates of BnOPR and AtOPR3, respectively. Fragmentation spectra were compared to databases and literature to unequivocally confirm the chemical structure. For Arabidopsides, *m*/*z* 263.166 [*dn*-*cis*-OPDA + H^+^]^+^ and *m*/*z* 291.197 [*cis*-OPDA + H^+^]^+^ were considered characteristic fragments for the cyclopentenone moieties. The fragmentation spectra revealed that both *cis*-OPDA and/or *dn*-*cis*-OPDA moieties (in Arabidopsides A and B) are reduced to either 3-oxo-2-(2-pentenyl)-cyclopentane-1-octanoic acid (OPC-8) or 3-oxo-2-(2-pentenyl)-cyclopentane-1-hexanoic acid (OPC-6), respectively, leading to a 4.014 Da mass difference between substrate and product. Several markers, namely OPC-6 + glycerol + hexose, OPC-8 + glycerol + hexose, *cis*-OPDA + glycerol, and *cis*-OPDA + glycerol + hexose, were identified by fragment spectra. Based on their retention time and further observed markers with the same retention time, they could be derived from in-source fragmentation of Arabidopsides. However, the abundance of OPC-6+glycerol+hexose within AtOPR1 and AtOPR2 containing reactions could also indicate these are different metabolites within the metabolite extract. A full list of markers is provided in Dataset S2. Altogether, the *ex vivo* approach confirmed the broad substrate spectrum of the tested OYEs. Nevertheless, differences between the substrate preferences were visible and confirmed the closer relationship of BnOPR to AtOPR3 than to AtOPR1 and AtOPR2.

**Figure 6.**
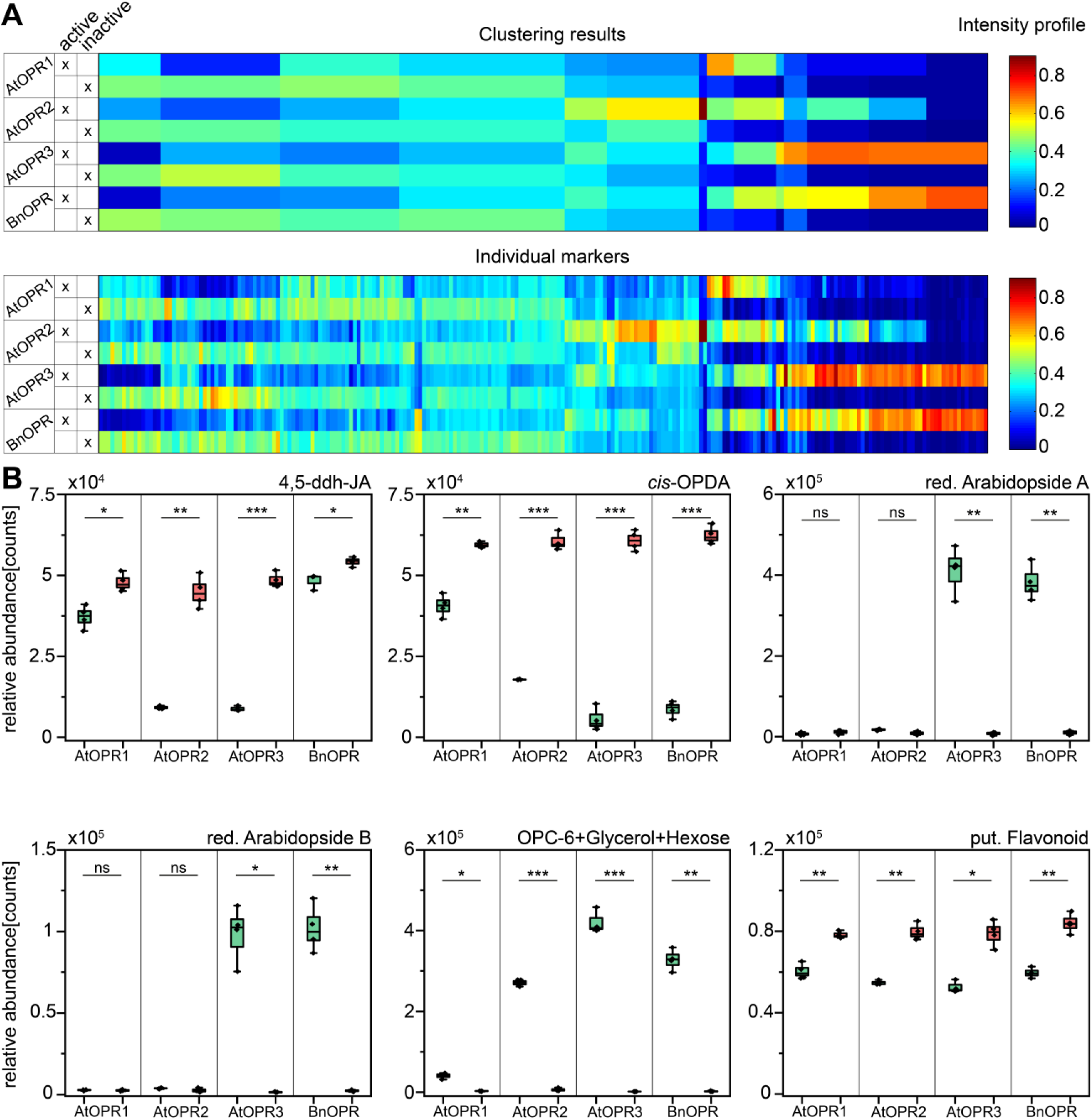
*Ex vivo* assay and relative intensity of selected markers. MTBE two-phase extracts of wounded leaves of *Atopr2 Atopr3* were incubated with heterologously expressed and purified AtOPR1, AtOPR2, AtOPR3, and BnOPR. 1 mM NADPH was added to prevent cofactor shortage. Reactions were stopped after 120 min with 50 % (*v/v*) MeOH and analyzed via UPLC-HRMS. Controls were conducted with heat-inactivated enzyme. Data were processed with Profinder for peak picking and peak alignment (cut-off 4000 counts). Features were ranked and filtered by ANOVA coupled to multiple testing by Benjamini-Hochberg with a false discovery rate of 0.001 with MarVis Filter. **A** Features were clustered by mean in a one-dimensional self-organizing map (1D-SOM) with MarVis Cluster. **B** Selected markers were confirmed by MS/MS fragmentation experiments. Boxplots show the relative abundance of respective markers. Green boxes represent the amounts in samples with active enzymes and red boxes represent samples with heat-inactivated enzymes. *P*-values were calculated with the Student’s *t*-test and marked accordingly: \**P*<0.01, ** *P* <0.001,*** *P* <0.0001. Each reaction was conducted four times. **Alt text:** Figures A and B present the results of the *ex vivo* metabolomic assay of AtOPR1 to 3 and BnOPR. Figure A shows one-dimensional self-organizing maps. The top panel shows the clustering results, and the lower panel the individual markers within this clustering. To the left, the respective enzymes and whether the enzyme was active or inactive are stated for each row. Figure B shows selected markers with substrate or product patterns for the respective enzymes. The displayed boxplots show each active sample in green and the inactive samples in red, sorted according to the tested enzymes. The identities of each boxplot from left to right and top to bottom are: 4,5-didehydro-jasmonic acid, reduced Arabidopside B, *cis*-OPDA, OPC-6+Glycerol+Hexose, reduced Arabidopside A, and a putative flavonoid.

### *Brevibacillus* spp. have a growth promoting effect on *P. patens*

The origin of the bacterial contamination in the *P. patens* genome remains unsolved but most probably stems from a co-cultivation of the strain that was grown for the initial V1 assembly (Rensing *et al*., 2008). During decontamination of the V1 assembly, a complete Bacillus genome was assembled and removed (Lang *et al*., 2018). During decontamination of the V3 assembly, a few remaining bacterial scaffolds were removed, among them the one harbouring BnOPR (Bi *et al*., 2024). Of note, *P. physcomitrellae* was isolated from a *P. patens* lab culture in China (Zhou *et al*., 2015). This pedigree, as well as the one sequenced for the genome assembly, goes back to the same single spore isolate from UK (Haas *et al*., 2020). Such co-cultivated bacteria might mitigate growth-promoting effects. To test a potential role of *B. nitrificans* in the *P. patens* rizosphere itself, a close relative, *B. brevis*, and *P. physcomitrellae* were tested in growth assays and revealed growth-promoting effects on plant growth. *B. nitrificans* inoculated plants grew significantly bigger within 25 days, however, the effect appears to be weak (Figure 7). *B. brevis* inoculated gametophores, on the other hand, showed approximately double rhizoid length and slightly longer gametophores but reduced gametophore numbers per plant compared to controls (Figures S6+S7). *P. physcomitrellae* inoculation similarly led to increased gametophore length but did not alter gametophore numbers per plant. This could indicate a beneficial effect of *Brevibacillus spp.* within the *P. patens* rhizosphere, which could therefore explain the co-isolation of *B. nitrificans*.

**Figure 7.**
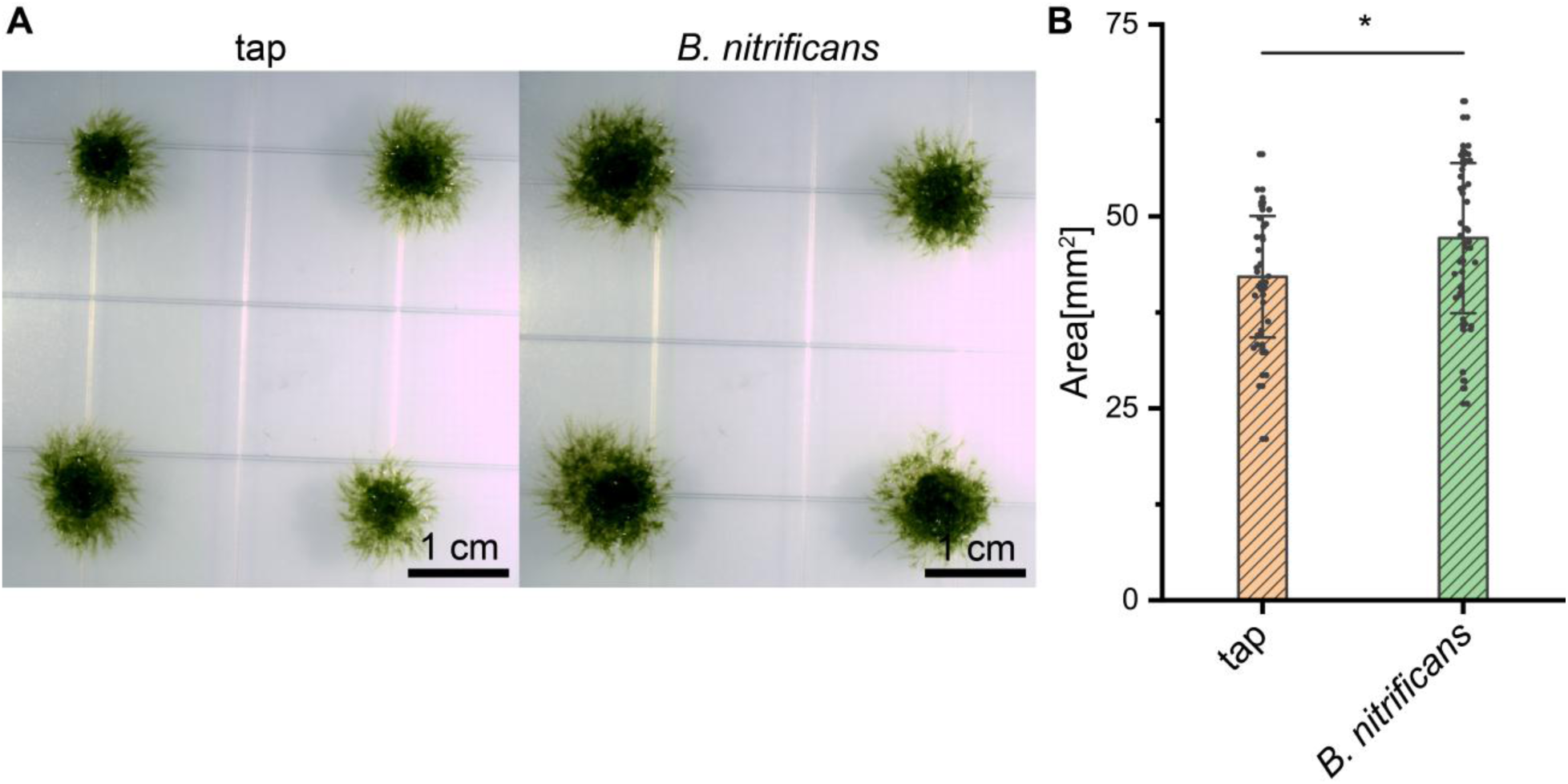
*B. nitrificans* promotes growth in *P. patens.* **A** *P. patens* gametophores were cultivated for 25 days under long-day conditions on solid minimal medium supplemented with 5 % (*w/v*) sucrose. Each colony was inoculated with 20 μL solution of *B. nitrificans* at 25,000 cells/mL. Pictures show representative results for colony growth for *B. nitrificans* and tap water inoculated. **B** Surface area was measured and compared in three independent experiments and for more than 35 plants per replicate (Mean ± SD, *P*<0.05, Student’s *t*-test). Scale bar: 1 cm. **Alt text:** Figures A and B show the growth experiments for *P. patens* incubated with tap water or *B. nitrificans* and the quantified colony area. Figure A shows four representative plants for the tap water control to the left and co-cultivated with *B. nitrificans* to the right. Figure B shows the quantified colony area in mm². Here, the *B. nitrificans* colonies are significantly larger compared to the tap water control.

The results shown in this publication demonstrate the functionality of BnOPR under *in vitro* and *in planta* conditions. The contamination of the *P. patens* genome occurred probably through co-isolation and sequencing of the bacterial genome. This is supported by the growth-promoting effects of *B. nitrificans* and *B. brevis*. Phylogenetic analysis revealed the bacterial origin of BnOPR and makes its occurrence as a result of horizontal gene transfer unlikely. Surprisingly, the functionality as an OPR was confirmed with *in planta* complementation experiments, which were supported with *in vitro* assays and a conclusive *ex vivo* approach. This could give implications for an ancient bacterial OYE with OPR-like functionality.

## Discussion

Although OYEs were discovered almost 100 years ago, their physiological role and their *in vivo* substrates remain mostly enigmatic (Böhmer *et al*., 2023; Warburg and Christian, 1932). The intensive *in vitro* characterization of OYEs, however, led to a still increasing number of substrates, reaction mechanisms, and consequently biotechnological applications (Scholtissek *et al*., 2017; Shi *et al*., 2020). The best studied group of OYEs with a confirmed physiological function is plant OPRs and their role in JA biosynthesis (Schaller and Stintzi, 2009). Here, we aim to expand our knowledge on this group of OYE by describing a putative OPR from the facultative anaerobic gram-positive bacterium *B. nitrificans*. The sequence was first identified in a phylogenetic analysis as a *P. patens* gene in genome V3 before decontamination. The occurrence of the *BnOPR* sequence in the *P. patens* genome could be the result of a co-isolation of both organisms. *Bacillus* species have been found to be associated with *P. patens* and a whole bacillus genome was amplified and had to be removed during the assembly of P. patens genome V1 (Rensing *et al*., 2008; Zhou *et al*., 2015). When tested for a growth-promoting effect of *B. brevis,* however, it could be demonstrated that gametophores and rhizoids are elongated when both species are grown together (Figures S6+S7). *B. nitrificans* shares a 99.4 % DNA similarity with *B. brevis* (Takebe *et al*., 2012). *B. nitrificans* also demonstrated a growth-promoting effect, but to a lesser extent (Figure 7). These results confirm earlier observations of the growth-promoting effect of *B. brevis* on cotton (*Gossypium hirsutum*) and canola (de Freitas *et al*., 1997; Nehra *et al*., 2016). Further research demonstrated a potential application of *B. brevis* as a biocontrol organism against *Fusarium* spp. (Chandel *et al*., 2010; Joo *et al*., 2015).

The enzymatic characterization of BnOPR showed that it is able to reduce *cis*-OPDA not only *in vitro* but as well *in vivo* as demonstrated in complementation experiments with the male infertile *A. thaliana* mutant *Atopr3* (Stintzi and Browse, 2000). The expression under the control of the *CaMV35S* promoter and the localization to the peroxisome is sufficient to restore silique formation and increase JA levels by 3-5 fold, compared to the mutant, although the complementing lines do not reach WT levels (Figure 4). This observation is supported by recent experiments, in which the double mutant *Atopr2 Atopr3* was not only complemented with AtOPR3, but also with AtOPR1 and AtOPR2 (Mekkaoui *et al*., 2025). In this study, the complementing lines also could not restore WT JA levels, but formed siliques in a WT-like manner. Consequently, the threshold of JA signaling in the flowers appears to be low compared to the actual amount of JA produced in WT flowers.

To compare the enzymatic properties of BnOPR with AtOPR3, a substrate library was tested with both recombinant enzymes. Under these conditions, AtOPR3 reduced *cis*-OPDA, 2-cyclopenten-1-one, 3-methyl-2-cyclopenten-1-one and 4,5-ddh-JA. These findings are in line with previous studies (Maynard *et al*., 2019; Mekkaoui *et al*., 2025; Schaller *et al*., 2000). The reported KM values for AtOPR3 were 35 µM and 28.3 µM for *cis*-OPDA, respectively and the KM value of BnOPR for *cis*-OPDA determined as 53 µM within this study was in a similar concentration range (Table 1). AtOPR3 showed for 8-*iso*-Prostaglandin A1 a lower KM of 22 µM and a turnover number of 0.23 s^-1^ (Han *et al*., 2011). As assessed by *in vitro* assays, prostaglandin A2 was an efficient substrate for BnOPR, even though apparent kinetic parameters were not determined. The second best tested substrate based on the KM value was prednisone, which was a better substrate for BnOPR compared to AtOPR3. These results indicate a preference of BnOPR for bulkier substrates, which is in line with the active site structure and the surface models (Figures 2 and S3). The *ex vivo* assay also revealed that Arabidopsides were reduced by AtOPR3 and BnOPR, but not to a lesser extent by AtOPR1 and AtOPR2 (Figure 6). This is to our very best knowledge the first report showing the capability of OPRs to reduce the cyclopentenone moieties in Arabidopsides and may suggest for a function in the wound response of wounded leaves. Upon wound-induced tissue disruption AtOPR3 may have direct access to Arabidosides leading to a faster reduction and thereby detoxification of *cis*-OPDA moieties before they may be released from the galactolipid backbone.

Many OYEs are very promiscuous and accept substrates ranging from cyclic and aromatic α,β-unsaturated enones to aliphatic enones and nitro compounds (Stott *et al*., 1993; Vaz *et al*., 1995). Even bulkier molecules like sterols and α,β-unsaturated lipid-derived compounds, like Arabidopsides, can act as substrates. Surprisingly, OYE from *S. pasteurianum* was able to reduce members of most of these groups, but not prednisone (Vaz *et al*., 1995). The same enzymes, SpOYE1 and SpOYE2, were also able to reduce *cis*-OPDA in *in vitro* assays (Schaller and Weiler, 1997). Therefore, it can be speculated that OYE1 and OYE2 would also be able to complement the *Atopr3* male infertile phenotype.Taking this into account, it highlights the importance of characterizing members of different classes of OYEs to exploit the full potential of this enzyme family for their physiological functions and efficient biocatalytic transformations. Previous attempts to predict substrate specificity were aimed at the active site residues. Especially, Tyr74/Tyr242 for AtOPR1, Tyr76/Tyr244 for AtOPR2, and Phe74/His245 were discussed as the determining residues for stereoselectivity of OPDA isomers (Figure 2). Even if the stereoselectivity was not addressed in this study, the Asn72 and Tyr233 residues at the respective positions in BnOPR differ from the described residues, while still achieving reduction of *cis*-OPDA (Breithaupt *et al*., 2009). It has to be mentioned that these two amino acids are just predictied based on modeling, while a X-ray structure is lacking to pinpoint exactly the residues for substrate specificity. BLAST analysis revealed that Tyr371 from AtOPR3 is missing in the BnOPR structure (Figure S2C), leading to an overall open active site (Figure S2). This could allow even bulkier molecules to bind and access the FMN cofactor. The increased space for substrate binding may have a drawback in substrate specificity.

Earlier studies revealed the involvement of AtOPRs in the detoxification of explosives by reduction of the reactive nitro-groups (Beynon *et al*., 2009). BnOPR, on the other hand, was not able to reduce 2,4-DNP or 3,5-dinitrosalicylic acid. Therefore, it appears that nitro compounds are not within the substrate scope of BnOPR.

The results of this study presented a bacterial *OYE* that was wrongfully assigned as a *P. patens* gene, to be a functional OPR in complementation lines. *Ex vivo* and *in vitro* assays support the activity in an AtOPR3-like manner with an affinity towards bulkier substrates. Further, the results indicate that *Brevibacillus* spp. are located in proximity to *P. patens* rhizoids, with potentially beneficial effects on plant growth, which led to the co-isolation and sequencing of this contaminating sequence. A new genome version did not show this sequence (Bi *et al*., 2024).

## Supplementary data

The following supplementary data are available at JXB online.

**Dataset S1.** Species list maximum-likelihood phylogeny.

**Dataset S2.** *Ex vivo* metabolome markers.

**File S1.** Labeled phylogenetic tree.

## Author contribution

MK, KF, SR and IF experimental design and conceptualization; MK, EH, LP, AK, LB, MH and LS performed the experiments; MK, EH, KF, LB and LS conducted data analysis and processing; MK and IF wrote the manuscript. Every author proof read and accepted the manuscript.

## Acknowledgements and funding statement

We thank Sabine Freitag for her help with the synthesis and purification of *cis*-OPDA. We thank Susanne Mester and Andrea Nickel for their help with plant cultivation. This research has been funded by the Deutsche Forschungsgemeinschaft (DFG; GRK 2172: 273134146, 118910695, 495720893, ZUK 45/2010, SPP 2237: 528076711) to I. F. and J. d. V.. M. K., L. P. and A.K. acknowledge the support by the PRoTECT program of the Goettingen Graduate Center for Neuroscience, Biophysics and Molecular Biosciences (GGNB).

## Data availability

All primary data of UHPLC-HRMS are available at MetaboLights.

## Abbreviations

1D-SOM: one-dimensional self-organizing map
HRMS: high-resolution mass spectrometry
JA: Jasmonic acid
MeJA: Methyl jasmonate
MR: morphine reductase
NerA: glycerol trinitrate reductase
*cis*-OPDA: *cis*-12-*oxo*-phytodienoic acid
OPR: *cis*-12-*oxo*-phytodienoic acid reductase
OYE: Old Yellow Enzyme
PETNR: pentaerythritol tetranitrate reductase
SEC: size exclusion chromatography
TNT: 2,4,6-trinitrotoluene
TOYE: thermophilic “ene” reductase
UHPLC: ultra-high-pressure-liquid-chromatography

## Supplementary material

**Figure S1.**
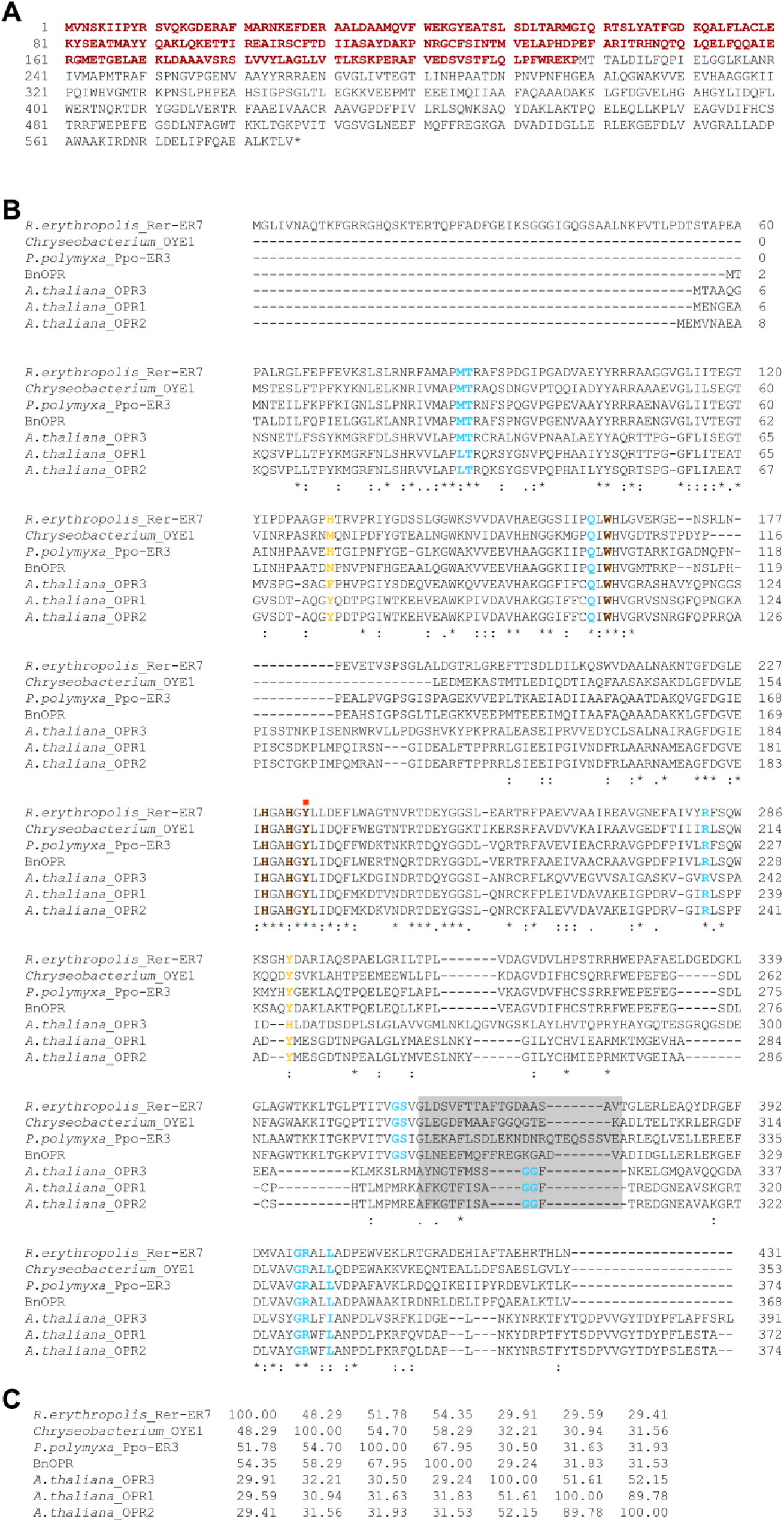
Multiple sequence alignment of BnOPR, AtOPRs, and selected Class IV OYEs. **A** Sequence of annotated Pp3s58_150 gene product. TetR7AcR sequence is highlighted in red. **B** Sequences were subjected to multiple sequence alignment using Clustal Omega (Madeira *et al*., 2024). Residues important for substrate binding and catalysis are colored brown, with the two residues determining substrate specificity in OPRs marked in orange and residues interacting with the FMN cofactor marked in light blue. Grey box marks the loop extension, characteristic for Class IV OYEs. The tyrosine residue important for catalysis is marked with a red square. **C** Percent identity matrix of the selected sequences.

**Figure S2.**
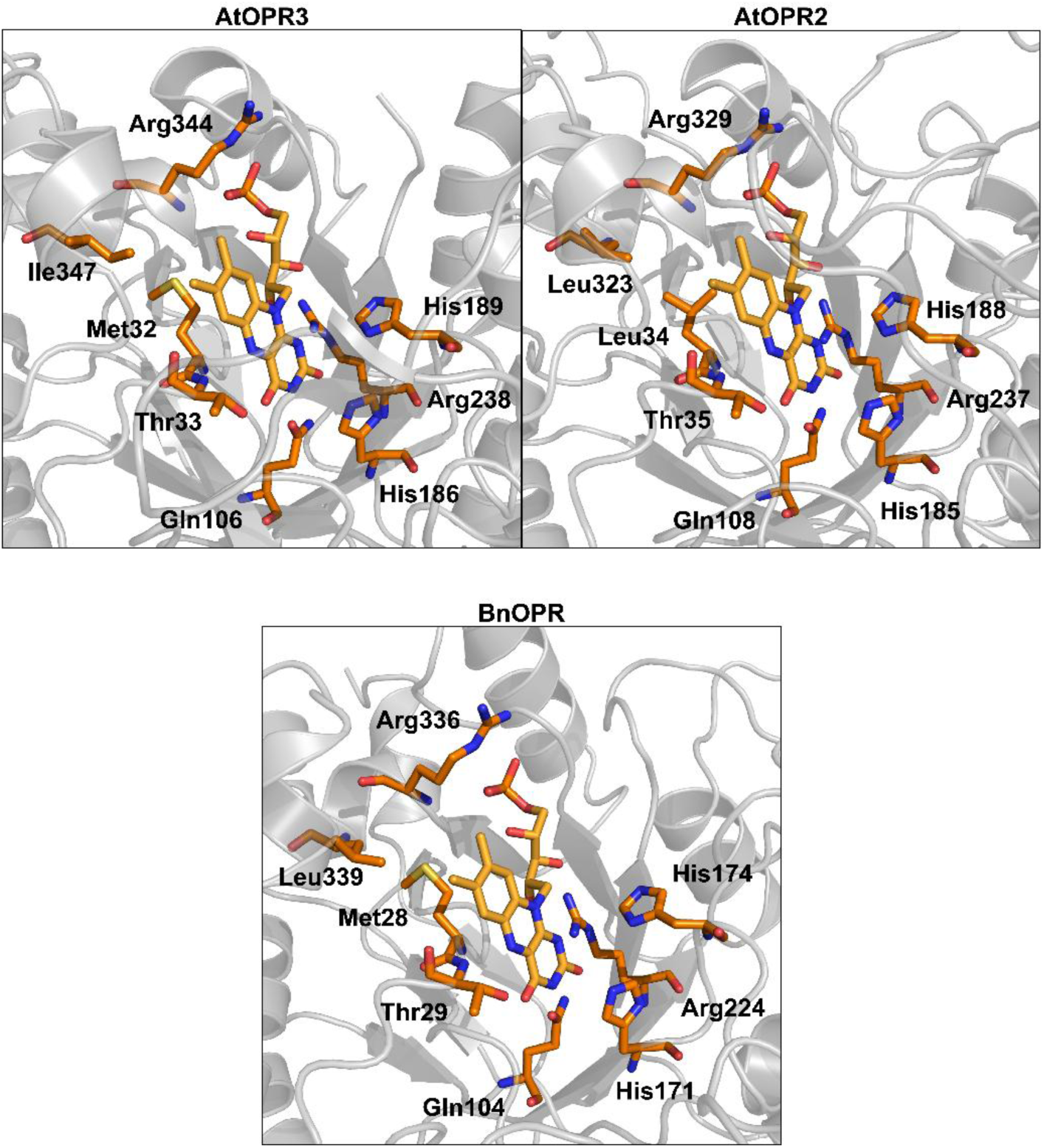
FMN binding site of AtOPR2, AtOPR3 and BnOPR. FMN binding sites of AtOPR3 (PDB: 2G5W) and Alphafold2 models of AtOPR2 and BnOPR were visualized according to multiple sequence alignment. Residues interacting with the FMN cofactor are shown in orange sticks with respective residue numbers.

**Figure S3.**
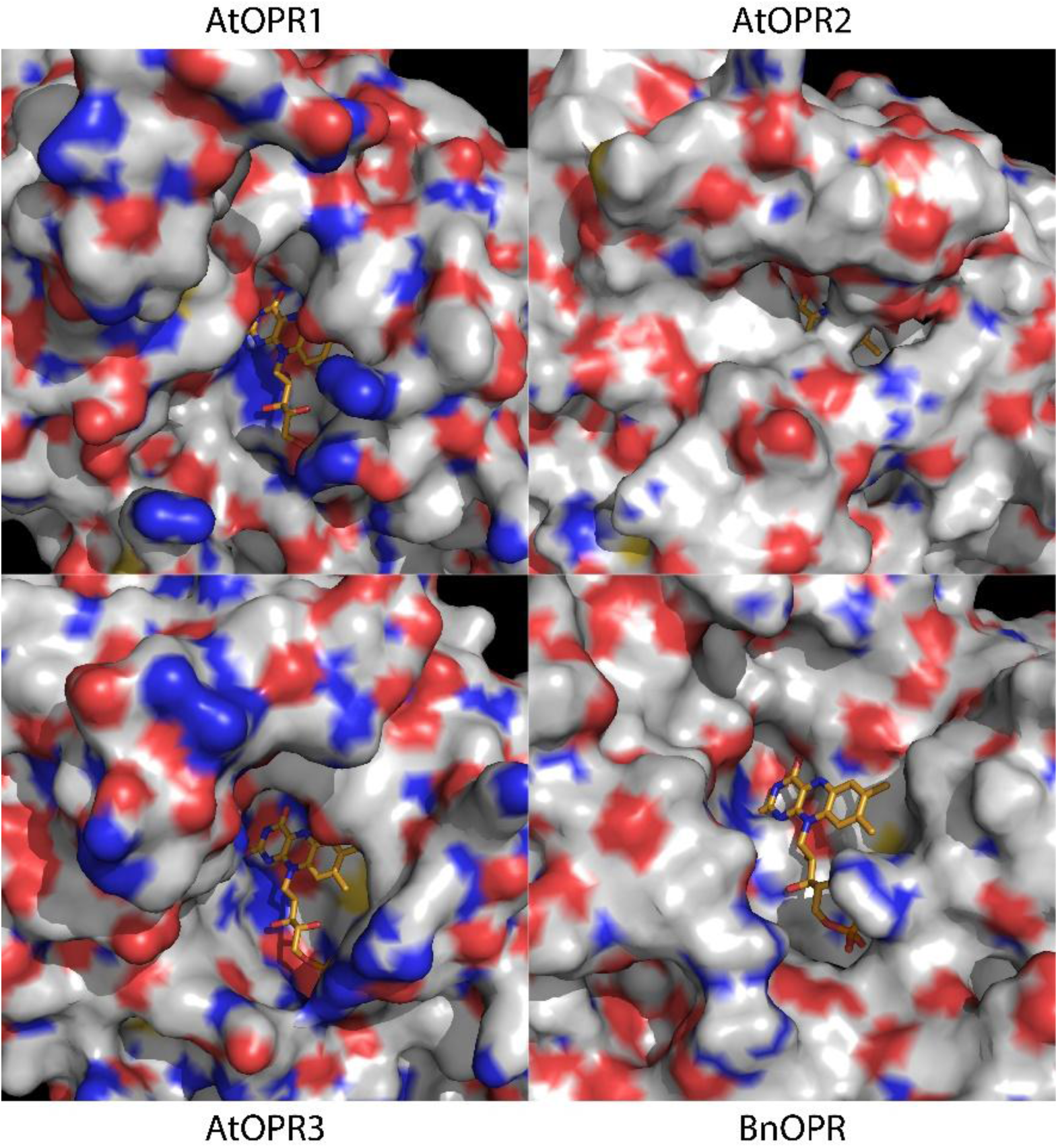
Surface model of the active sites of BnOPR and AtOPR1-3. Comparison of active site/ binding pocket from the crystal structures of AtOPR1 (PDB: 1VJI) and AtOPR3 (PDB: 2G5W). BnOPR and AtOPR2 were modeled using Alphafold2 (Mirdita *et al*., 2022). Panels show the enzyme surface with oxygen in red, nitrogen in blue and sulfur atoms in yellow. FMN was superimposed into the models from AtOPR3 crystal structure.

**Figure S4.**
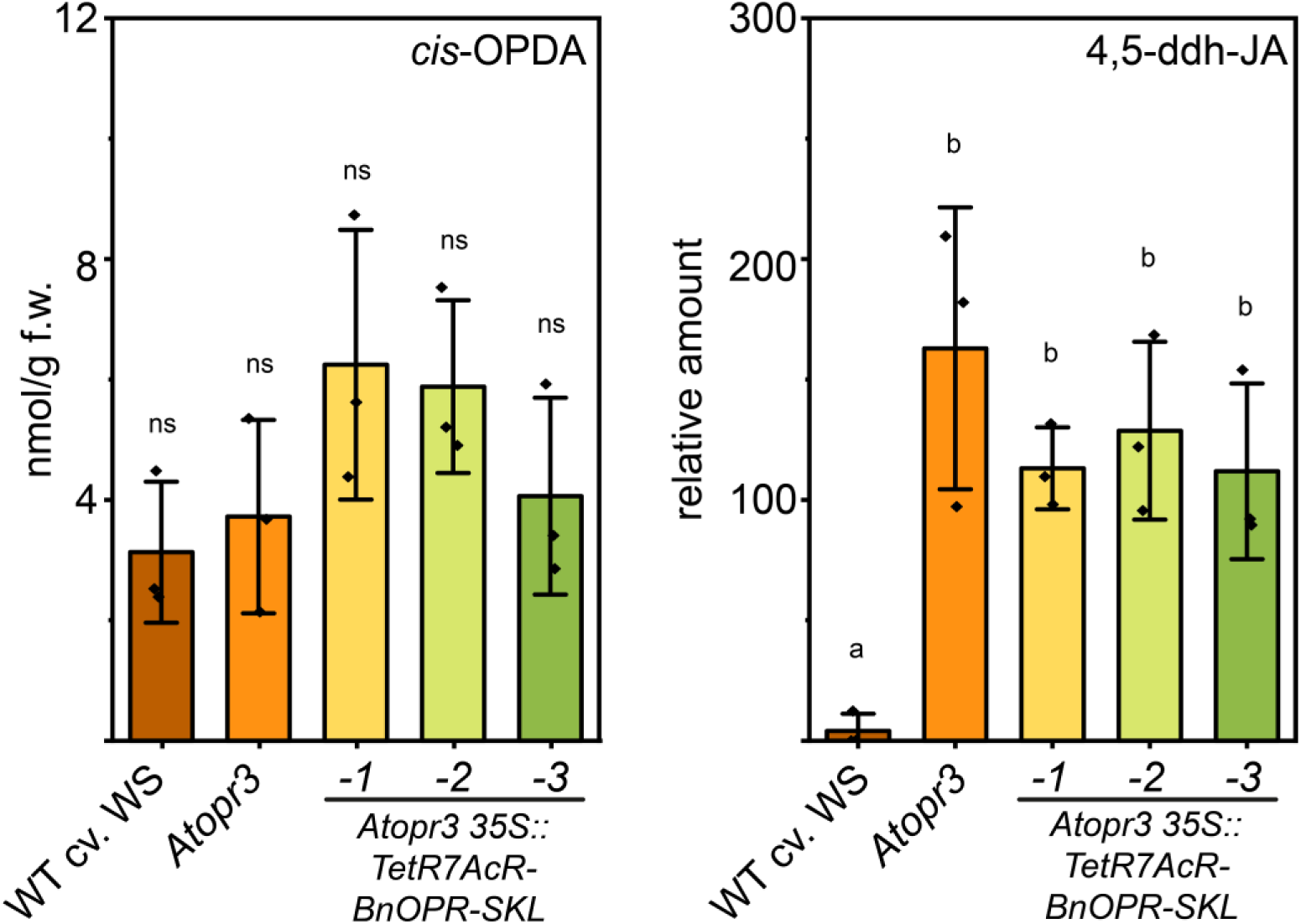
*cis*-OPDA and 4,5-ddh-JA levels in complementing *Atopr3* lines. *cis*-OPDA and 4,5-ddh-JA were measured in MeJA treated and wounded leaves after one hour. Samples were analyzed as described before (Herrfurth and Feussner, 2020). Data was acquired from three biological replicates. Statistics were calculated by one-way ANOVA followed by *post-hoc* Tukey’s HSD test and are indicated by different letters (*P*<0.05).

**Figure S5.**
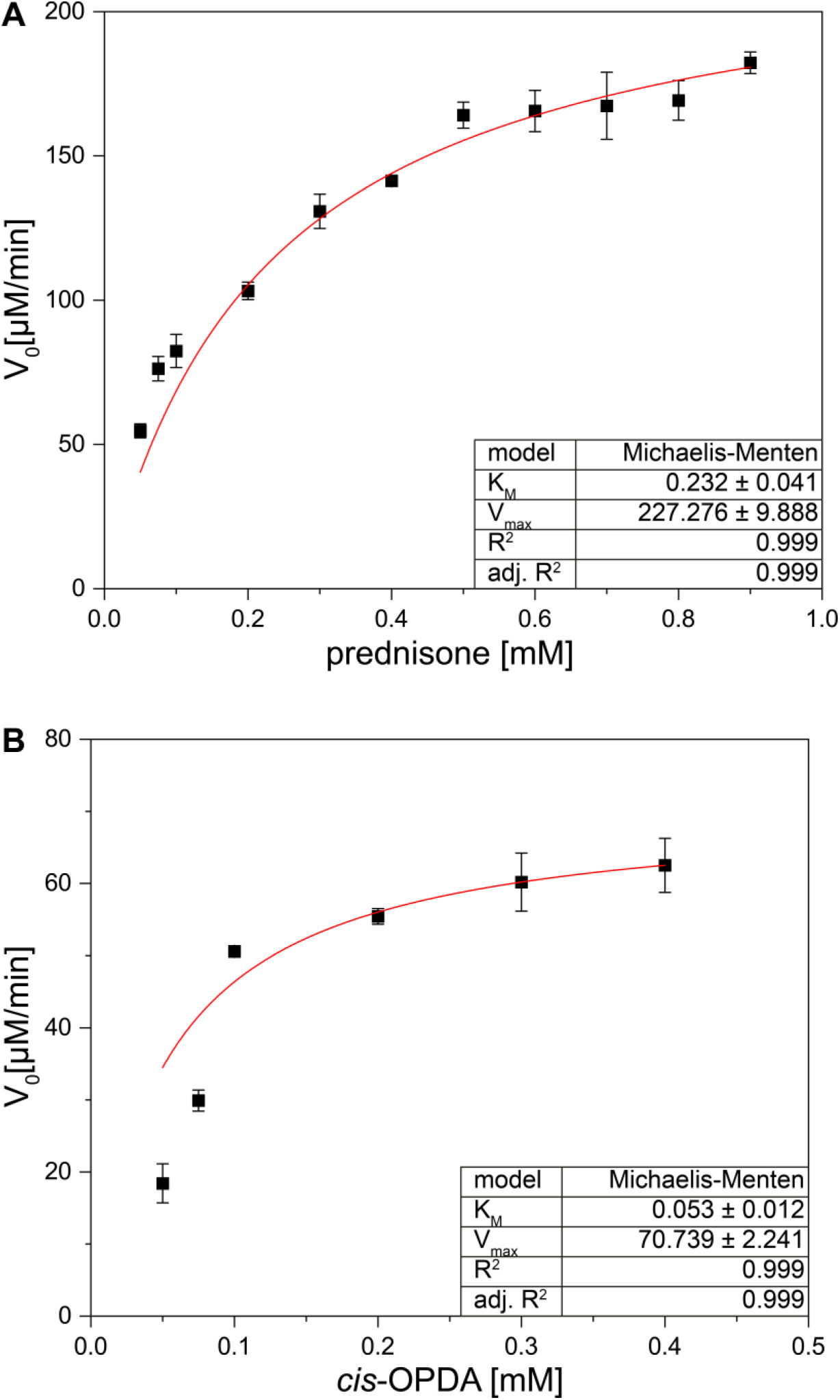
Michaelis-Menten fits of prednisone and *cis*-OPDA kinetic measurements. Heterologously expressed and purified BnOPR was subjected to kinetic measurements with NADPH and altering concentrations of prednisone or *cis*-OPDA. Reaction was set up in 20 mM HEPES (pH 7.5) and was incubated at 25 °C. Absorbance change at 340 nm was observed using a Thermo Genesys 50, plotted and fitted with Origin. Results of three measurements were fitted according to Michaelis-Menten equation. Kinetic parameters were determined by the software and are shown in the tables on the right of the graphs.

**Figure S6.**
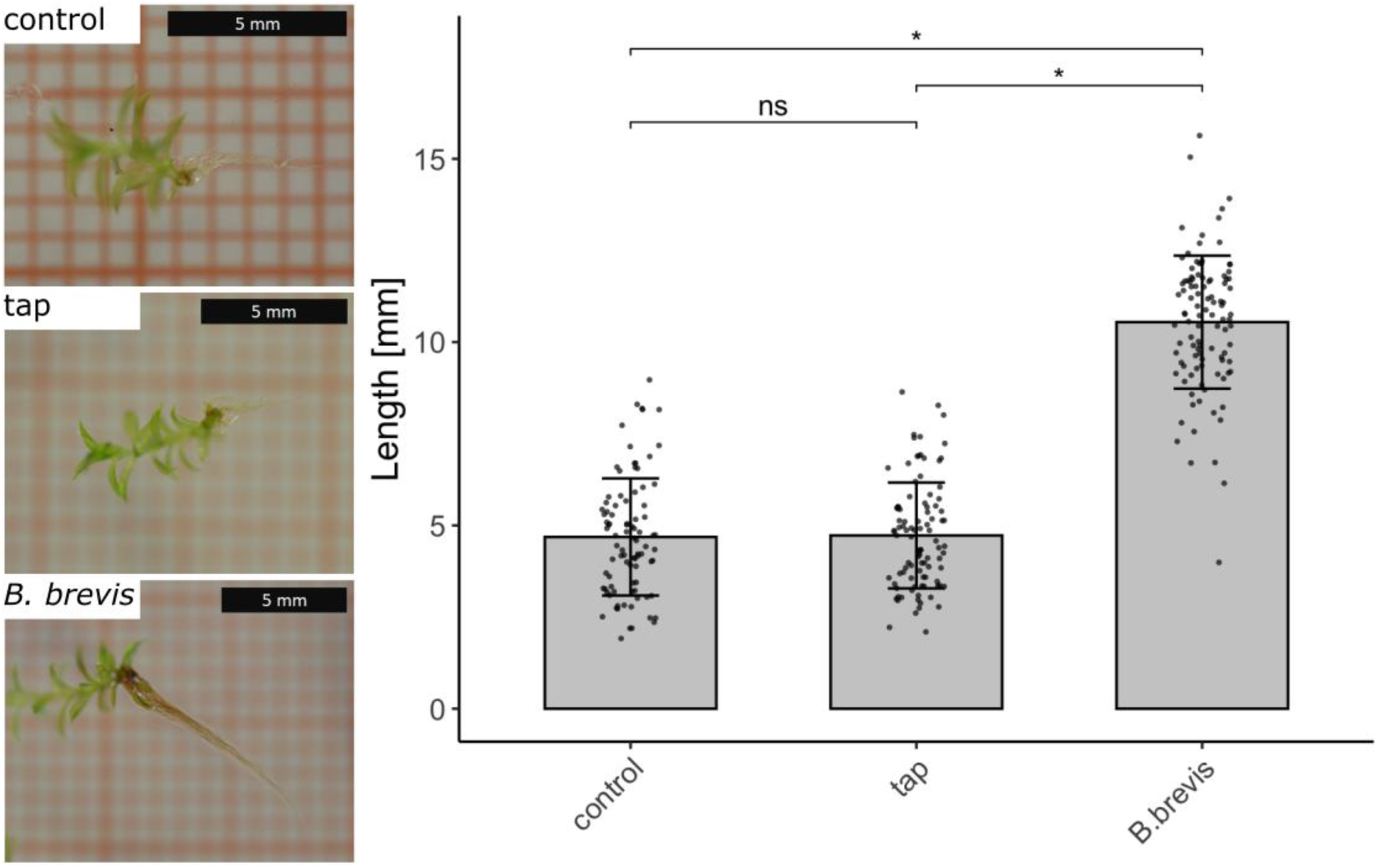
*B. brevis* promotes growth of rhizoids in *P. patens*. *P. patens* gametophores were cultivated for 25 days under long-day conditions on M-medium supplemented with 5 g/L sucrose. Each gametophore was inoculated with 20 μL solution of *B. brevis* at 25,000 cells/mL. Pictures show representative results for rhizoid growth in non-inoculated, tap water and *B. brevis* inoculated treatment. Rhizoid length was measured and compared in three independent experiments and for at least 26 plants per replicate (Mean ± SD, *P*<0.001, Student’s *t*-test).

**Figure S7.**
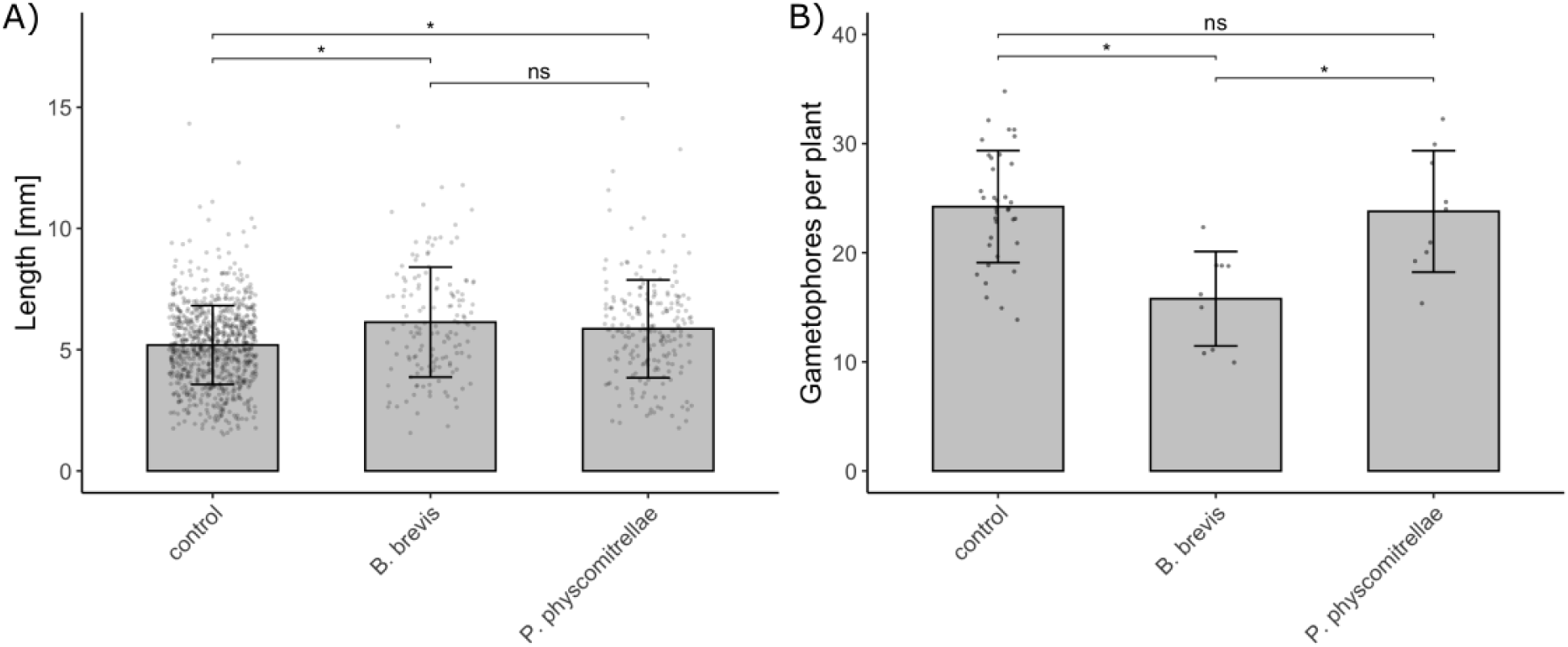
*B. brevis* and *P. physcomitrellae* alter gametophore growth in *P. patens*. *P. patens* cultures were spotted on medium and precultured for four weeks before inoculation. **A** *P. patens* shows elongated gametophores when inoculated with *B. brevis* or *P. physcomitrellae*. Gametophore length was measured in three independent experiments with at least three plants. **B** *P. patens* shows reduced gametophore numbers per plant when inoculated with *B. brevis*. Gametophore numbers were counted in three independent experiments on at least three plants (Mean ± SD, P<0.05, Student’s t-test).

---

**Table S1.**
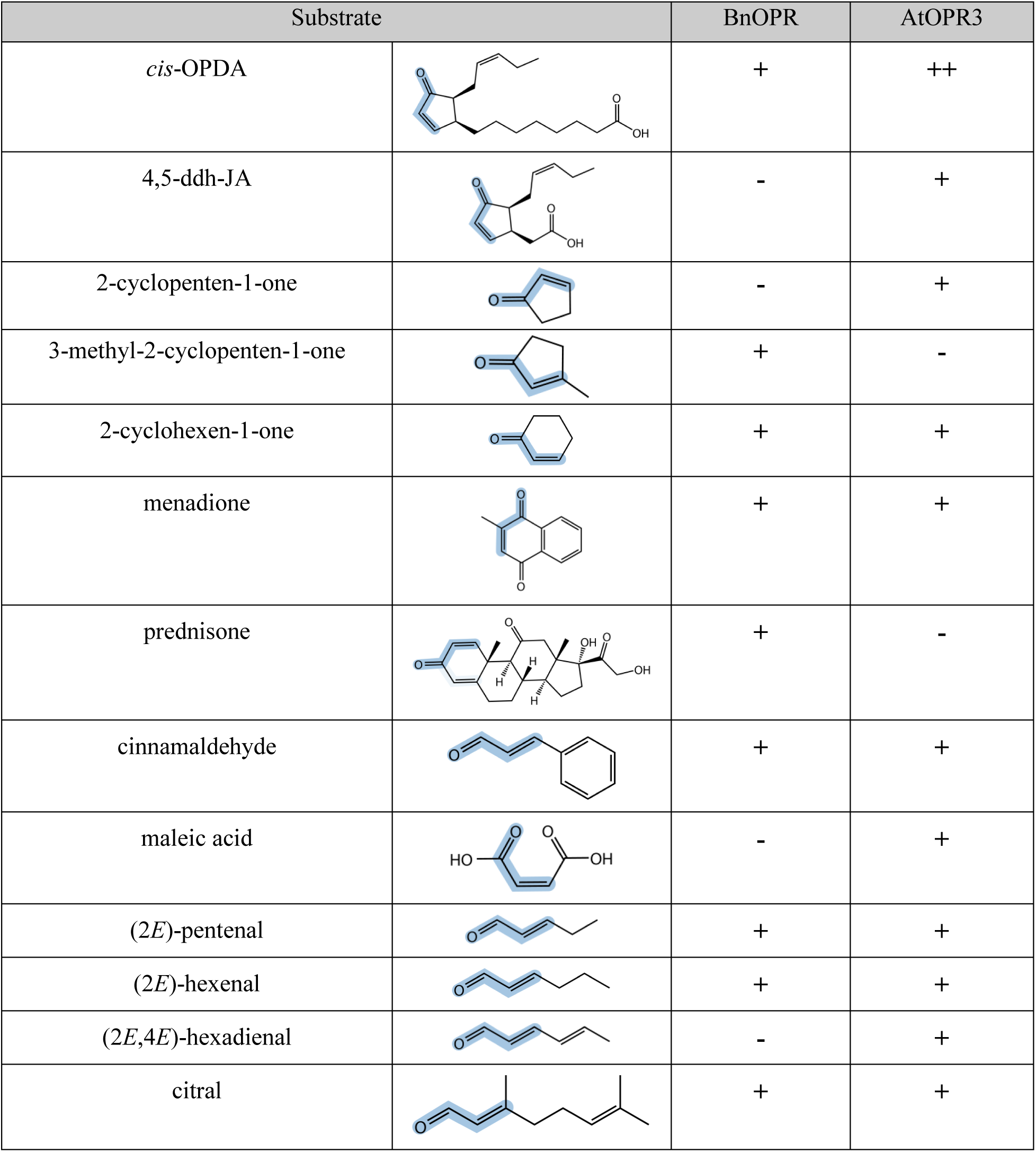
Qualitative comparison of BnOPR with AtOPR3. To compare the substrate preferences of BnOPR and AtOPR3, enzymes were tested in assays, consisting of the respective substrate and NADPH. Reactions were incubated at 30°C and stopped after 10 min. The reactions were analyzed by UHPLC-HRMS for substrate consumption and product formation. No significant reduction activity is marked (-), while a nearly complete conversion is marked (++). Everything in between is marked (+). Each reaction was measured three times.

| Substrate |  | BnOPR | AtOPR3 |
| --- | --- | --- | --- |
| *cis*-OPDA |  | + | ++ |
| 4,5-ddh-JA |  | - | + |
| 2-cyclopenten-1-one |  | - | + |
| 3-methyl-2-cyclopenten-1-one |  | + | - |
| 2-cyclohexen-1-one |  | + | + |
| menadione |  | + | + |
| prednisone |  | + | - |
| cinnamaldehyde |  | + | + |
| maleic acid |  | - | + |
| (2*E*)-pentenal |  | + | + |
| (2*E*)-hexenal |  | + | + |
| (2*E*,4*E*)-hexadienal |  | - | + |
| citral |  | + | + |

## References

Adalbjörnsson BV, Toogood HS, Fryszkowska A, Pudney CR, Jowitt TA, Leys D, Scrutton NS. 2010. Biocatalysis with thermostable enzymes: Structure and properties of a thermophilic ‘ene’-reductase related to old yellow enzyme. Chembiochem 11, 197–207.

Barna T, Messiha HL, Petosa C, Bruce NC, Scrutton NS, Moody PCE. 2002. Crystal structure of bacterial morphinone reductase and properties of the C191A mutant enzyme. Journal of Biological Chemistry 277, 30976–30983.

Bécard G, Fortin JA. 1988. Early events of vesicular–arbuscular mycorrhiza formation on Ri T-DNA transformed roots. New Phytologist 108, 211–218.

Beynon ER, Symons ZC, Jackson RG, Lorenz A, Rylott EL, Bruce NC. 2009. The role of oxophytodienoate reductases in the detoxification of the explosive 2,4,6-trinitrotoluene by Arabidopsis. Plant Physiology 151, 253–261.

Bi G, Zhao S, Yao J, Wang H, Zhao M, Sun Y, Hou X, Haas FB, Varshney D, Prigge M, Rensing SA, Jiao Y, Ma Y, Yan J, Dai J. 2024. Near telomere-to-telomere genome of the model plant *Physcomitrium patens*. Nature Plants 10, 327–343.

Böhmer S, Marx C, Goss R, Gilbert M, Sasso S, Happe T, Hemschemeier A. 2023. Chlamydomonas reinhardtii mutants deficient for Old Yellow Enzyme 3 exhibit increased photooxidative stress. Plant Direct 7, e480.

Breithaupt C, Strassner J, Breitinger U, Huber R, Macheroux P, Schaller A, Clausen T. 2001. X-ray structure of 12-oxophytodienoate reductase 1 provides structural insight into substrate binding and specificity within the family of OYE. Structure 9, 419–429.

Breithaupt C, Kurzbauer R, Lilie H, Schaller A, Strassner J, Huber R, Macheroux P, Clausen T. 2006. Crystal structure of 12-oxophytodienoate reductase 3 from tomato: Self-inhibition by dimerization. Proceedings of the Nationtal Academy of Sciences USA 103, 14337–14342.

Breithaupt C, Kurzbauer R, Schaller F, Stintzi A, Schaller A, Huber R, Macheroux P, Clausen T. 2009. Structural basis of substrate specificity of plant 12-oxophytodienoate reductases. Journal of Molecular Biology 392, 1266–1277.

Chandel S, Allan EJ, Woodward S. 2010. Biological control of *Fusarium oxysporum* f.sp. *lycopersici* on tomato by *Brevibacillus brevis*. Journal of Phytopathology 158, 470–478.

Chehab EW, Kim S, Savchenko T, Kliebenstein D, Dehesh K, Braam J. 2011. Intronic T-DNA insertion renders Arabidopsis *opr3* a conditional jasmonic acid-producing mutant. Plant Physiology 156, 770–778.

Chini A, Monte I, Zamarreño AM, Hamberg M, Lassueur S, Reymond P, Weiss S, Stintzi A, Schaller A, Porzel A, García-Mina JM, Solano R. 2018. An OPR3-independent pathway uses 4,5-didehydrojasmonate for jasmonate synthesis. Nature Chemical Biology 14, 171–178.

Clough SJ, Bent AF. 1998. Floral dip: a simplified method for *Agrobacterium*-mediated transformation of *Arabidopsis thaliana*. The Plant Journal 16, 735–743.

de Freitas JR, Banerjee MR, Germida JJ. 1997. Phosphate-solubilizing rhizobacteria enhance the growth and yield but not phosphorus uptake of canola (*Brassica napus L*.). Biology and Fertility of Soils 24, 358–364.

Egener T, Granado J, Guitton MC, Hohe A, Holtorf H, Lucht JM, Rensing SA, Schlink K, Schulte J, Schween G, Zimmermann S, Duwenig E, Rak B, Reski R. 2002. High frequency of phenotypic deviations in Physcomitrella patens plants transformed with a gene-disruption library. BMC Plant Biology 2, 6.

Feussner K, Abreu IN, Klein M, Feussner I. 2023. Chapter Twelve - Metabolite fingerprinting: A powerful metabolomics approach for marker identification and functional gene annotation. In: Jez J, ed. Methods in Enzymology, Vol. 680. Cambridge, MA: Academic Press, 325–350.

Fitzpatrick TB, Amrhein N, Macheroux P. 2003. Characterization of YqjM, an old yellow ezyme homolog from *Bacillus subtilis* involved in the oxidative stress response. Journal of Biological Chemistry 278, 19891–19897.

Fox BG, Malone TE, Johnson KA, Madson SE, Aceti D, Bingman CA, Blommel PG, Buchan B, Burns B, Cao J, Cornilescu C, Doreleijers J, Ellefson J, Frederick R, Geetha H, Hruby D, Jeon WB, Kimball T, Kunert J, Markley JL, Newman C, Olson A, Peterson FC, Phillips Jr. GN, Primm J, Ramirez B, Rosenberg NS, Runnels M, Seder K, Shaw J, Smith DW, Sreenath H, Song J, Sussman MR, Thao S, Troestler D, Tyler E, Tyler R, Ulrich E, Vinarov D, Vojtik F, Volkman BF, Wesenberg G, Wrobel RL, Zhang J, Zhao Q, Zolnai Z. 2005. X-ray structure of Arabidopsis At1g77680, 12-oxophytodienoate reductase isoform 1. Proteins: Structure, Function, and Bioinformatics 61, 206–208.

Fox KM, Karplus PA. 1994. Old yellow enzyme at 2 A resolution: overall structure, ligand binding, and comparison with related flavoproteins. Structure 2, 1089–1105.

French CE, Nicklin S, Bruce NC. 1996. Sequence and properties of pentaerythritol tetranitrate reductase from *Enterobacter cloacae* PB2. Journal of Bacteriology 178, 6623–6627.

French CE, Nicklin S, Bruce NC. 1998. Aerobic degradation of 2,4,6-trinitrotoluene by Enterobacter cloacae PB2 and by pentaerythritol tetranitrate reductase. Applied and Environmental Microbiology 64, 2864–2868.

French CE, Rosser SJ, Davies GJ, Nicklin S, Bruce NC. 1999. Biodegradation of explosives by transgenic plants expressing pentaerythritol tetranitrate reductase. Nature Biotechnology 17, 491–494.

Goetz S, Hellwege A, Stenzel I, Kutter C, Hauptmann V, Forner S, McCaig B, Hause G, Miersch O, Wasternack C, Hause B. 2012. Role of cis-12-oxo-phytodienoic acid in tomato embryo development. Plant Physiology 158, 1715–1727.

Haas FB, Fernandez-Pozo N, Meyberg R, Perroud P-F, Göttig M, Stingl N, Saint-Marcoux D, Langdale JA, Rensing SA. 2020. Single nucleotide polymorphism charting of *P. patens* reveals accumulation of somatic mutations during *in vitro* culture on the scale of natural variation by selfing. Frontiers in Plant Science 11, 813.

Han BW, Malone TE, Kim DJ, Bingman CA, Kim H-J, Fox BG, Phillips Jr. GN. 2011. Crystal structure of *Arabidopsis thaliana* 12-oxophytodienoate reductase isoform 3 in complex with 8-*iso* prostaglandin A1. *Proteins: Structure*, Function, and Bioinformatics 79, 3236–3241.

Heilmann M, Iven T, Ahmann K, Hornung E, Stymne S, Feussner I. 2012. Production of wax esters in plant seed oils by oleosomal cotargeting of biosynthetic enzymes. Journal of Lipid Research 53, 2153–2161.

Herrfurth C, Feussner I. 2020. Quantitative jasmonate profiling using a high-throughput UPLC-NanoESI-MS/MS method. In: Champion A, Laplaze L, eds. Jasmonate in Plant Biology: Methods and Protocols, Vol. 2085. New York, NY: Springer US, 169–187.

Howe GA, Major IT, Koo AJ. 2018. Modularity in jasmonate signaling for multistress resilience. Annual Review of Plant Biology 69, 387–416.

Joo HJ, Kim H-Y, Kim L-H, Lee S, Ryu J-G, Lee T. 2015. A *Brevibacillus* sp. antagonistic to mycotoxigenic *Fusarium* spp. Biological Control 87, 64–70.

Kaever A, Lingner T, Feussner K, Göbel C, Feussner I, Meinicke P. 2009. MarVis: a tool for clustering and visualization of metabolic biomarkers. BMC Bioinformatics 10, 92.

Kalyaanamoorthy S, Minh BQ, Wong TK, Von Haeseler A, Jermiin LS. 2017. ModelFinder: fast model selection for accurate phylogenetic estimates. Nature Methods 14, 587–589.

Katoh K, Standley DM. 2013. MAFFT multiple sequence alignment software version 7: improvements in performance and usability. Molecular Biology and Evolution 30, 772–780.

Kunz CF, Goldbecker ES, de Vries J. 2025. Functional genomic perspectives on plant terrestrialization. Trends in Genetics 41, 617–629.

Lang D, Ullrich KK, Murat F, Fuchs J, Jenkins J, Haas FB, Piednoel M, Gundlach H, Van Bel M, Meyberg R, Vives C, Morata J, Symeonidi A, Hiss M, Muchero W, Kamisugi Y, Saleh O, Blanc G, Decker EL, van Gessel N, Grimwood J, Hayes RD, Graham SW, Gunter LE, McDaniel S, Hoernstein SNW, Larsson A, Li F-W, Perroud P-F, Phillips J, Ranjan P, Rokshar DS, Rothfels CJ, Schneider L, Shu S, Stevenson DW, Thümmler F, Tillich M, Villarreal A JC, Widiez T, Wong GK-S, Wymore A, Zhang Y, Zimmer AD, Quatrano RS, Mayer KFX, Goodstein D, Casacuberta JM, Vandepoele K, Reski R, Cuming AC, Tuskan J, Maumus F, Salse J, Schmutz J, Rensing SA. 2018. The *Physcomitrella patens* chromosome-scale assembly reveals moss genome structure and evolution. The Plant Journal 93, 515–533.

Letunic I, Bork P. 2024. Interactive Tree of Life (iTOL) v6: recent updates to the phylogenetic tree display and annotation tool. Nucleic Acids Res 52, W78–W82.

Li W, Liu B, Yu L, Feng D, Wang H, Wang J. 2009. Phylogenetic analysis, structural evolution and functional divergence of the 12-oxo-phytodienoate acid reductase gene family in plants. BMC Evolutionary Biology 9, 90.

Madeira F, Madhusoodanan N, Lee J, Eusebi A, Niewielska A, Tivey ARN, Lopez R, Butcher S. 2024. The EMBL-EBI Job Dispatcher sequence analysis tools framework in 2024. Nucleic Acids Research 52, W521–W525.

Maynard D, Kumar V, Sproß J, Dietz K-J. 2019. 12-oxophytodienoic acid reductase 3 (OPR3) functions as NADPH-dependent α,β-ketoalkene reductase in detoxification and monodehydroascorbate reductase in redox homeostasis. bioRxiv, 820381.

Mekkaoui K, Baral R, Smith F, Klein M, Feussner I, Hause B. 2025. Transcriptomics and trans-organellar complementation reveal limited signaling of 12-*cis*-oxo-phytodienoic acid during early wound response in *Arabidopsis*. Nature Communications 16, 6684.

Mirdita M, SchC, Moriwaki Y, Heo L, Ovchinnikov S, Steinegger M. 2022. ColabFold: making protein folding accessible to all. Nature Methods 19, 679–682.

Mohnike L, Rekhter D, Huang W, Feussner K, Tian H, Herrfurth C, Zhang Y, Feussner I. 2021. The glycosyltransferase UGT76B1 modulates *N*-hydroxy-pipecolic acid homeostasis and plant immunity. The Plant Cell 33, 735–749.

Nehra V, Saharan BS, Choudhary M. 2016. Evaluation of *Brevibacillus brevis* as a potential plant growth promoting rhizobacteria for cotton (*Gossypium hirsutum*) crop. SpringerPlus 5, 948.

Nguyen L-T, Schmidt HA, Von Haeseler A, Minh BQ. 2015. IQ-TREE: a fast and effective stochastic algorithm for estimating maximum-likelihood phylogenies. Molecular Biology and Evolution 32, 268–274.

Ni B, Feussner K. 2023. Chapter Eleven - Ex vivo metabolomics—A hypothesis-free approach to identify native substrate(s) and product(s) of orphan enzymes. In: Jez J, ed. Methods in Enzymology, Vol. 680. Cambridge, MA: Academic Press, 303–323.

Niino YS, Chakraborty S, Brown BJ, Massey V. 1995. A new old yellow enzyme of *Saccharomyces cerevisiae*. Journal of Biological Chemistry 270, 1983–1991.

Oberdorfer G, Binter A, Wallner S, Durchschein K, Hall Ml, Faber K, Macheroux P, Gruber K. 2013. The structure of glycerol trinitrate reductase NerA from *Agrobacterium radiobacter* reveals the molecular reason for nitro- and ene-reductase activity in OYE homologues. Chembiochem 14, 836–845.

Pak H, Wang H, Kim Y, Song U, Tu M, Wu D, Jiang L. 2021. Creation of male-sterile lines that can be restored to fertility by exogenous methyl jasmonate for the establishment of a two-line system for the hybrid production of rice (*Oryza sativa* L.). Plant Biotechnology Journal 19, 365–374.

Peters C, Buller R. 2019. Linear enzyme cascade for the production of (-)-*iso*-isopulegol. Zeitschrift für Naturforschung C 74, 63–70.

Peters C, Frasson D, Sievers M, Buller R. 2019. Novel old yellow enzyme subclasses. Chembiochem 20, 1569–1577.

Rensing SA, Lang D, Zimmer AD, Terry A, Salamov A, Shapiro H, Nishiyama T, Perroud P-F, Lindquist EA, Kamisugi Y, Tanahashi T, Sakakibara K, Fujita T, Oishi K, Shin-I T, Kuroki Y, Toyoda A, Suzuki Y, Hashimoto S-i, Yamaguchi K, Sugano S, Kohara Y, Fujiyama A, Anterola A, Aoki S, Ashton N, Barbazuk WB, Barker E, Bennetzen JL, Blankenship R, Cho SH, Dutcher SK, Estelle M, Fawcett JA, Gundlach H, Hanada K, Heyl A, Hicks KA, Hughes J, Lohr M, Mayer K, Melkozernov A, Murata T, Nelson DR, Pils B, Prigge M, Reiss B, Renner T, Rombauts S, Rushton PJ, Sanderfoot A, Schween G, Shiu S-H, Stueber K, Theodoulou FL, Tu H, Van de Peer Y, Verrier PJ, Waters E, Wood A, Yang L, Cove D, Cuming AC, Hasebe M, Lucas S, Mishler BD, Reski R, Grigoriev IV, Quatrano RS, Boore JL. 2008. The *Physcomitrella* genome reveals evolutionary insights into the conquest of land by plants. Science 319, 64–69.

Robescu MS, Cendron L, Bacchin A, Wagner K, Reiter T, Janicki I, Merusic K, Illek M, Aleotti M, Bergantino E, Hall M. 2022. Asymmetric proton transfer catalysis by stereocomplementary old yellow enzymes for C═C bond isomerization reaction. ACS Catalysis 12, 7396–7405.

Saito K, Thiele DJ, Davio M, Lockridge O, Massey V. 1991. The cloning and expression of a gene encoding old yellow enzyme from *Saccharomyces carlsbergensis*. Journal of Biological Chemistry 266, 20720–20724.

Schaller A, Stintzi A. 2009. Enzymes in jasmonate biosynthesis - Structure, function, regulation. Phytochemistry 70, 1532–1538.

Schaller F, Weiler EW. 1997. Molecular cloning and characterization of 12-oxophytodienoate reductase, an enzyme of the octadecanoid signaling pathway from *Arabidopsis thaliana* - Structural and functional relationship to yeast old yellow enzyme. Journal of Biological Chemistry 272, 28066–28072.

Schaller F, Hennig P, Weiler EW. 1998. 12-Oxophytodienoate-10,11-reductase: occurrence of two isoenzymes of different specificity against stereoisomers of 12-oxophytodienoic acid. Plant Physiology 118, 1345–1351.

Schaller F, Biesgen C, Mussig C, Altmann T, Weiler EW. 2000. 12-Oxophytodienoate reductase 3 (OPR3) is the isoenzyme involved in jasmonate biosynthesis. Planta 210, 979–984.

Scholtissek A, Ullrich SR, Mc, SchlC6mann M, Paul CE, Tischler D. 2017. A thermophilic-like ene-reductase originating from an acidophilic iron oxidizer. Applied Microbiology and Biotechnology 101, 609–619.

Shi Q, Wang H, Liu J, Li S, Guo J, Li H, Jia X, Huo H, Zheng Z, You S, Qin B. 2020. Old yellow enzymes: structures and structure-guided engineering for stereocomplementary bioreduction. Applied Microbiology and Biotechnology 104, 8155–8170.

Stintzi A, Browse J. 2000. The Arabidopsis male-sterile mutant, opr3, lacks the 12-oxophytodienoic acid reductase required for jasmonate synthesis. Proceedings of the National Academy of Sciences of the United States of America 97, 10625–10630.

Stintzi A, Weber H, Reymond P, Browse J, Farmer EE. 2001. Plant defense in the absence of jasmonic acid: The role of cyclopentenones. Proceedings of the Nationtal Academy of Sciences USA 98, 12837–12842.

Stott K, Saito K, Thiele DJ, Massey V. 1993. Old yellow enzyme. The discovery of multiple isozymes and a family of related proteins. Journal of Biological Chemistry 268, 6097–6106.

Strassner J, Schaller F, Frick UB, Howe GA, Weiler EW, Amrhein N, Macheroux P, Schaller A. 2002. Characterization and cDNA-microarray expression analysis of 12-oxophytodienoate reductases reveals differential roles for octadecanoid biosynthesis in the local versus the systemic wound response. The Plant Journal 32, 585–601.

Takebe F, Hirota K, Nodasaka Y, Yumoto I. 2012. *Brevibacillus nitrificans* sp. nov., a nitrifying bacterium isolated from a microbiological agent for enhancing microbial digestion in sewage treatment tanks. International Journal of Systematic and Evolutionary Microbiology 62, 2121–2126.

Tatusova T, Ciufo S, Fedorov B, O’Neill K, Tolstoy I. 2013. RefSeq microbial genomes database: new representation and annotation strategy. Nucleic Acids Research 42, D553–D559.

Tischler D, Gädke E, Eggerichs D, Gomez Baraibar A, Mügge C, Scholtissek A, Paul CE. 2020. Asymmetric reduction of (*R*)-carvone through a thermostable and organic-solvent-tolerant ene-reductase. Chembiochem 21, 1217–1225.

Toogood HS, Gardiner JM, Scrutton NS. 2010. Biocatalytic reductions and chemical versatility of the old yellow enzyme family of flavoprotein oxidoreductases. ChemCatChem 2, 892–914.

Vaz ADN, Chakraborty S, Massey V. 1995. Old yellow enzyme: Aromatization of cyclic enones and the mechanism of a novel dismutation reaction. Biochemistry 34, 4246–4256.

Vick BA, Zimmerman DC. 1986. Characterization of 12-oxo-phytodienoic acid reductase in corn. Plant Physiology 80, 202–205.

Warburg O, Christian W. 1932. Über das neue Oxydationsferment. Naturwissenschaften 20, 980–981.

Wasternack C, Feussner I. 2018. The oxylipin pathways: Biochemistry and function. Annual Review of Plant Biology 69, 363–386.

White DW, Frkic RL, Iamurri S, Keshavarz-Joud P, Blue T, Jackson CJ, Copp J, Lutz S. 2025. Expanding the biocatalytic and oxidative landscape of the old yellow enzyme family. Protein Science 34, e70363.

Yan Y, Christensen S, Isakeit T, Engelberth J, Meeley R, Hayward A, Emery RJN, Kolomiets MV. 2012. Disruption of OPR7 and OPR8 reveals the versatile functions of jasmonic acid in maize development and defense. The Plant Cell 24, 1420–1436.

Zhou X, Nan Guo G, Qi Wang L, Lan Bai S, Li Li C, Yu R, Hong Li Y. 2015. *Paenibacillus physcomitrellae* sp. nov., isolated from the moss *Physcomitrella patens*. International Journal of Systematic and Evolutionary Microbiology 65, 3400–3406.

